# A survey of *cis* regulatory non-coding RNA involved in bacterial virulence

**DOI:** 10.1101/2021.11.03.467129

**Authors:** Mohammad Reza Naghdi, Samia Djerroud, Katia Smail, Jonathan Perreault

## Abstract

Study of pathogenesis in bacteria is important to find new drug targets to treat bacterial infections. Pathogenic bacteria, including opportunists, express numerous so-called virulence genes to escape the host natural defenses and immune system. Regulation of virulence genes is often required for bacteria to infect their host. Such regulation can be achieved by *cis*-regulatory RNAs, like the metabolite-binding riboswitches or thermoregulators. In spite of the hundreds of RNA families annotated as cis-regulatory, there are relatively few examples of non-coding RNAs (ncRNAs) in 5′-UnTranslated Regions (UTRs) of bacteria described to regulate downstream virulence genes. To reassess the potential roles of such regulatory elements in bacterial pathogenesis, we collected genes important for virulence from different databases and evaluated the presence of ncRNAs in their UTRs to highlight the potential role of this type of gene regulation for virulence and, at the same time, get insight on some of the physical and chemical triggers of virulence.

## 1 Introduction

Virulence factors (VFs) are in large part responsible of the relative pathogenicity of bacteria and often counter the host immune system (Casadevall and Pirofski, 2003). The expression of VFs is controlled in accordance to environmental cues and signaling changes from the host (DiRita et al., 2000), or changes like temperature increase while moving from water to a mammalian host, or signals of microbiota living in the host, to give only a few examples (Bäumler and Sperandio, 2016) The communication between the host and microorganisms is known as inter-kingdom signaling and is crucial for activation of VFs which activates cascades of signaling pathways (Hughes and Sperandio, 2008). For example, it has been shown that sensing a specific metabolite such as fucose in intestines is crucial for effective colonization (Pacheco et al., 2012). The importance of two component regulatory systems has also been shown, like PhoPQ in Mg^2+^ sensing (García Véscovi et al., 1996, Véscovi et al., 1997) or QseC, a quorum-sensing regulator which detects eukaryote hormones for VF regulation (Clarke et al., 2006, Rasko et al., 2008). Other examples of physicochemical triggers include pH variation and quorum sensing that control expression of streptococcal pyrogenic exotoxin B (SpeB), a VF in *Streptococcus pyogenes* (Do et al., 2019).

Tight regulation of VFs improves the ability of pathogens to infect their host (Caldelari et al., 2013). Quick regulation is a key for successful pathogenicity, which is expected in a highly changing environment like hosts, especially when the latter react to the invasion (Fris and Murphy, 2016).

The regulation with RNA has been shown to be more effective than regulation with proteins in some contexts (Gripenland et al., 2010). Noncoding RNAs (ncRNAs) are a heterogeneous group of RNA that do not code for proteins but instead directly enact a function, often related to gene control. Regulation carried out by ncRNAs can impact one or several genes during transcription or translation (Eddy, 2001). The ncRNAs can be divided into two major groups as *cis*-regulatory or non *cis*-regulatory *(trans)* RNAs. *Cis-*regulatory RNAs are located mostly in the 5’-UnTranslated Regions (UTRs) of genes and have a direct effect on expression of the downstream gene. Regulation can occur at the transcriptional or translational level (Abduljalil, 2018). Good examples of *cis-*regulatory elements are riboswitches (Nahvi et al., 2002), RNA thermosensors (Morita et al., 1999) and T-boxes (Grundy and Henkin, 1993). For example, it has been shown that several VFs in *Listeria monocytogenes* are controlled by *cis-*regulatory RNAs such as riboswitches and thermoregulator RNAs (Lebreton and Cossart, 2017). The latter regulate the *prfA* gene (Johansson et al., 2002), a major regulator of virulence in *L. monocytogenes* (Leimeister-Wächterchter et al., 1990, Mengaud et al., 1991). *L. monocytogenes* also uses different flavors of a SAM riboswitch to regulate its virulence (Loh et al., 2009). The regulation of virulence genes *via* ncRNAs with a focus on sRNA was reviewed recently (Chakravarty and Massé, 2019). Here we evaluated *cis*-regulatory RNAs involved in regulation of virulence genes (VFs).

Increased use of RNAseq and improved bioinformatics prediction pipelines revealed numerous ncRNAs (Weinberg et al., 2010, Weinberg et al., 2017, Leonard et al., 2019, Stav et al., 2019) many of which we find may have an impact on virulence and may help expose more regulatory roles of ncRNAs *in vivo* (Hör et al., 2018). We used an exhaustive list of ncRNAs to explore their association to known virulence genes by combining data from different relational databases to provide a collection of *cis*-regulatory ncRNAs regulating VFs to assist research on virulence regulation.

## 2 Materials and Methods

### 2.1 Bioinformatics

In order to determine the existence of all *cis*-regulatory elements upstream of virulence genes, an exhaustive list of annotated VFs was established by using PATRIC database (Wattam et al., 2014) and two other databases: VFDB (Chen et al., 2016) and Victors (Sayers et al., 2019). The virulence is defined as a capacity of a bacteria to infect a host by using the VFs which help the bacteria colonize (Sharma et al., 2017) and escape the host immune system which results in infection and disease (Mao et al., 2015). However, VFs are also genes which do not affect the viability of bacteria outside their hosts (Brown et al., 2012).

Next, we looked for all *cis*-regulatory RNAs upstream of these genes by using the RiboGap database (Naghdi et al., 2017) and BLASTp (Altschul et al., 1990, Camacho et al., 2009). All the bacterial intergenic 5′-UTRs having ncRNAs (484,136 sequences) were extracted with their corresponding cds using RiboGap, i.e. all prokaryotic ncRNAs searched with cmsearch with Rfam’s covariance models, as well as a few more RNAs (Supplementary Data). BLASTp was then used to determine homology

between genes downstream of ncRNAs and the list of VFs (9019 genes). PERL scripts (supplementary data) were used to analyze the results obtained by BLASTp. To avoid getting genes with common domains, but that are non-orthologous, the BLASTp condition was set to 98% coverage for High Scoring Pair (HSP). The BLASTp result was then sorted to keep hits with at least 60% identity. Only *cis*-regulatory RNAs on the same strand as the downstream gene were taken into account, except for tRNAs (see below).

### 2.2 tRNA searches

The tRNAs were searched separately. RNA distance from start codon of VF was also taken in consideration. The search was carried out for the same virulence genes as described above. Because all genomes harbor many tRNAs, numerous genes are expected to have tRNAs upstream of their coding sequences just by chance, so samples of genes (three replicas of 100 randomly chosen genes) were also used to put results in context. To evaluate the presence of pseudo-tRNAs and also obtain information on tRNA identity, tRNAscan-SE (Chan and Lowe, 2019) was used instead of RiboGap annotations, but RiboGap was used to fetch all the UTRs.

### 2.2 Northern of co-transcribed tRNAs

Three tRNAs were identified upstream of “Elongation Factor Tu” (*Ef-Tu*) gene in *Neisseria*. To determine whether this gene is transcribed alone or co-transcribed with the tRNAs upstream, we selected *Neisseria gonorrhoeae, Neisseria meningitidis, Neisseria sicca* and *Neisseria elongata*. Oligonucleotides complementary to each of the three tRNAs upstream of the *EF-Tu* cds and to the cds itself were ordered from IDT to probe the membrane (Table S1). Similarly, an oligonucleotide complementary to the tmRNA for the four *Neisseria* species was ordered from IDT and used as control.

Northern blots were performed as previously described (Perreault et al., 2011). In brief, total RNA of *Neisseria gonorrhoeae, Neisseria meningitidis* MC58_NMB0124, *Neisseria sicca* and *Neisseria elongata* was migrated on a 6% polyacrylamide gel and then transferred onto nitrocellulose membrane (Amersham Hybond™ N^+^ from GE healthcare). The oligonucleotides were labeled in 5’ by using 5 pmoles of oligonucleotide, 2 µL ATP (γ-^32-^P), 1 µL of 10 U/µL polynucleotide T4 kinase and PNK buffer (NEB) in 20 µL, then incubated at 37°C for 1 h. The labeled products were then purified on denaturing 6% polyacrylamide gel. The labeled oligonucleotides were incubated with the membrane for 24 hours at 42° C in a rotating oven with hybridization buffer with SCC 5X prepared from SCC 20 X (175.3 g NaCl, 88.2 g sodium citrate in 1 L, pH 7.0) and the day after washed twice with SCC 2X, 1 % SDS and SCC 0.2 X, 0.1% SDS. Membranes were then exposed overnight on a phosphorimaging plate. The plate was scanned with a Typhoon FLA9500.

## 3 Results

### 3.1 *cis-*regulatory RNA distribution upstream of virulence factors

We decided to not limit our search to the genes listed as VFs in the PATRIC, Victors and VFBD databases. The focus of these databases is on experimentally validated VF genes and orthologs that follow stringent criteria (including similar genomic context), but can omit some orthologs in other pathogens. We even extended our survey of cis-regulatory ncRNAs to non-pathogens because regulation of a gene in such species can still be informative for their VF orthologs in pathogenic counterparts, or between different pathogens. For example, a gene encoding a magnesium-translocating P-type ATPase is regulated by the *ykoK* Mg^2+^ riboswitch in the non-pathogen *Lactococcus lactis* (Dann et al., 2007), by the Mg-sensor riboswitch in the pathogen *Salmonella enterica* (Cromie et al., 2006) or by the MgtC leader in *Klebsiella aerogenes* (Table 1 and S2 and S3) as well as the PhoP/PhoQ two component system (García Véscovi et al., 1996, Cromie et al., 2006), all indicative of a common regulatory signal, in spite of different mechanisms. Therefore, extending searches to VF orthologs may provide hints on the regulation of these genes in pathogens.

**Table 1.** Compilation of thermoregulators and riboswitches regulating VFs and homologs

| RNA | Hits | Genes | Most abundant virulence gene | Signal | VF hits |
| --- | --- | --- | --- | --- | --- |
| **Riboswitches** |  |  |  | **(metabolites)** |  |
| ydaO/yuaA leader | 159 | 2 | resuscitation-promoting factor RpfA | Cyclic di-AMP | 1 |
| Cyclic di-GMP-I | 6 | 1 | peptide chain release factor 3 | Cyclic di-GMP | 0 |
| FMN | 1 | 1 | catalase | FMN | 0 |
| TPP | 249 | 3 | Phosphomethyl pyrimidine kinase | TPP | 2 |
| ZMP/ZTP | 133 | 4 | Bifunctional phosphoribosyl amino imidazole carboxamide formyl transferase | ZMP/ZTP | 0 |
| Lysine | 182 | 5 | Aspartokinase | Lysine | 0 |
| Glycine | 76 | 3 | glycine dehydrogenase | Glycine | 0 |
| Purine | 777 | 3 | glutamine-hydrolyzing GMP synthase | Purine | 0 |
| SAM (S box) | 521 | 3 | ABC transporter substrate-binding protein | SAM | 1 |
| SAH | 2 | 1 | methionine synthase | SAH | 0 |
| Cobalamin | 154 | 3 | Glutaredoxin-like protein NrdH | Cobalamin | 1 |
| Fluoride | 3 | 1 | Type I glutamate ammonia ligase |  | 0 |
| Moco | 52 | 1 | Formate-tetrahydrofolate | molybdenum cofactor | 0 |
|  |  |  |  | **(cations)** |  |
| ykoK leader | 10 | 1 | PPE family protein | magnesium | 0 |
| Salmonella MgtC leader | 3443 | 6 | Mg<sup>2+</sup> transporter MgtC | magnesium | 2 |
| NiCo | 1 | 1 | ATP-dependent Clp protease ATP-binding subunit | nickel/cobalt | 1 |
| Listeria snRNA rli51 | 1068 | 3 | zinc metalloproteinase PrtA | (zinc) | 2 |
| **Thermoregulators** |  |  |  | **(temperature)** |  |
| cspA | 2624 | 5 | cold-shock protein | Temperature | 2 |
| PrfA UTR | 238 | 1 | listeriolysin transcriptional regulator PrfA | Temperature | 0 |
| ToxT 5' UTR | 2 | 1 | pilus/toxin transcriptional regulator ToxT | Temperature | 0 |
| CnfY 5' UTR | 6 | 1 | DUF4765 domain-containing protein | Temperature | 0 |
| RhlA 5' UTR ROSE like | 96 | 1 | rhamnosyltransferase 1 subunit A | Temperature | 0 |
| KatA 5' UTR | 39 | 1 | Catalase | Temperature | 0 |
| SodB 5' UTR | 360 | 2 | superoxide dismutase [Fe] | Temperature | 4 |
| SodC 5' UTR | 16 | 1 | superoxide dismutase | Temperature | 0 |
| LasI 5' UTR ROSE like | 26 | 1 | acyl-homoserine-lactone synthase | Temperature | 0 |
| shuA/chuA 5' UTR | 640 | 1 | TonB-dependent hemoglobin/transferrin/lactoferrin family receptor | Temperature | 2 |
| HtrA 5' UTR | 2847 | 4 | periplasmic serine endoprotease DegP | Temperature | 2 |
| TrxA 5' UTR | 1500 | 3 | thioredoxin TrxA | Temperature | 2 |
| LcrF | 144 | 3 | virulence regulon transcriptional activator | Temperature | 2 |
| AilA 5' UTR | 126 | 3 | attachment protein | Temperature | 0 |

RNA : name of RNA family (as in Rfam)
Hits: total number of times this RNA was found upstream of genes annotated as VFs or homologous to VFs.
Genes: Number of different genes (or protein encoded by those genes) annotated as VFs or homologous to VFs.
Most abundant virulence gene: Example of the corresponding gene (or protein encoded by those genes) that most frequently has this RNA upstream of it.
Signal: Signal detected by the RNA and that triggers the change in expression. For riboswitches (including for cations) the signal is a molecule; for thermoregulators, body temperature of homeotherm (warm-blooded) animals typically triggers virulence; (zinc) the *Listeria* snRNA rli51 is not a known riboswitch, but given the downstream gene, a possible connection with zinc is possible (as seen for the small RNA RhyB and iron for instance (Massé and Gottesman, 2002).
VF hits: hits corresponding only to genes annotated as VFs (and not homologs).

We found 95,943 genes associated with virulence (and orthologs) downstream of ncRNAs (Table 1 and Table S2). From these RNAs, we selected cis-regulatory RNAs (as annotated “type” in Rfam) based on the criteria described in materials and methods to produce compiled lists of RNA families already known to be *cis*-regulators (Table 1 and Tables S3, S4 and S5). This list includes 16 riboswitches for metabolites, 14 thermoregulatory RNAs and 4 cation-associated regulators (Table 1), as well as many additional ncRNAs such as the T-boxes, leucine-operon leader or PyrD leader (1,473 hits in the latter case) (Table S5). The purine riboswitch was found to be the most common riboswitch among *cis*-regulatory RNAs (777 instance), whereas FMN and NiCo were found just one time and many riboswitches, such as THF, guanidine (I, II and III) and fluoride riboswitches, were not observed with any genes associated with virulence. Among the cation associated ncRNAs, the most common RNA family is associated to a zinc metalloproteinase, *Listeria snRNA rli51*, followed by the Mg^2+^ riboswitch and *ykoK*. Thermoregulators are clearly important ncRNAs regarding pathogenesis as 14 families of such ncRNAs are found upstream of ∼9,000 instances of VFs (or VF homologs). The most abundant thermoregulator identified is HtrA 5’ UTR, followed by *cspA* followed by TrxA-5’UTR and shuA/chuA 5’ UTR. While this may appear as few, compared to the 35 RNA *cis-*thermoregulator families, most of the other families have relatively few representatives or are found only in taxons never associated with infections, such as *cyanobacteria*.

### 3.2 tRNA upstream of virulence factors

We observed tRNAs upstream of hundreds of genes (Table S6). Interestingly, several VFs have pseudo-tRNAs in their UTR, such as *clpP* encoding a protease; and numerous genes have tRNA sequences on the opposite polarity in their 5′-UTR, like the *rnr* encoded ribonuclease for many *Betaproteobacteria* species (Table S7).

In numerous cases, tRNAs are found close enough to the downstream gene to suggest co-transcription (Tables S6 and S8). We tried to determine whether *Ef-Tu* was co-transcribed with the three tRNAs observed upstream of its cds in several *Neisseria* strains. The tRNA closest to *Ef-Tu* is at only 46 bases from its start codon, leaving little room for a promoter. The northern blot result reveals the three tRNAs (Tyr, Gly, Thr) are co-transcribed (supplementary Figure S1 and S2), but co-transcription with *Ef-Tu* was not apparent. Nevertheless, other instances of co-transcription are very likely in 56 cases where tRNAs are less than 40 bases apart from the downstream VF (or VF homolog). In some cases, the tRNAs even overlap the annotated 5′ portion of the coding sequence, such as in several strains of *Helicobacter pylori* for a gene encoding an outer membrane protein. Given precedents of tRNA^Sec^ that overlap the coding sequence of the *selB* gene close to its 3’ end (Mukai, 2021), tRNAs overlapping the 5’ end to enact gene control are easy to envision.

### 3.3 Transcription Terminators

We evaluated existing “Rho-independent transcription terminators” (RiTTs, Table S9) and “Rho-dependent transcription terminators” (RTTs, Table S10) for VFs as we did for other ncRNAs. However, because transcription terminators are very common, we only evaluated those upstream of VFs (as annotated in PATRIC, VFDB or Victors). These results can recapitulate several instances of RiTT deemed responsible of riboregulation as determined by Term-seq (Dar et al., 2016) (Table S11 and supplementary information). For example, *rli51* is a cis-regulatory RNA in *L. monocytogenes* that we highlight as associated to VFs and where we predict a RiTT, which is corroborated by the Term-seq results of Dar et al. 2016. This illustrates the usefulness of RiboGap to rapidly gather information on intergenic sequences and infer hypotheses that can be evaluated. As for the tRNA analyses, we considered distance from the start codon, we found that 189 and 6,403 predicted RiTTs and RTTs, respectively, are at less than 40 bases from the start codon, suggesting some transcription regulation independent from the promoter.

## 4 Discussion

To our knowledge, the results obtained from RiboGap and virulence databases are the most exhaustive inventory to date for known VFs related to *cis*-regulatory RNAs. Several hits are already well known as ncRNAs that regulate virulence factors (eg. thermoregulator RNA and *prfA* gene (Johansson et al., 2002)). The importance of thermoregulator RNAs in virulence of *Yersinia* (Nuss et al., 2017) and other species like *Shigella*.sp and pathogenic *E. coli* (Heroven et al., 2017) has been discussed by others. The prevalence of thermoregulator RNAs is high, as this is an excellent way for bacteria that can live in soil or water to determine they have moved to a warm-blooded animal.

Similarly, it is known that metals play important roles in virulence (Papp-Wallace and Maguire, 2006, Broder et al., 2016, Guragain et al., 2016, Imazawa et al., 2016, Palmer and Skaar, 2016, Wedekind et al., 2017). It is thus also not surprising that the list includes thousands of ncRNAs which appear connected to metal cations, in one way or another. The role of Mg^2+^ in virulence was previously connected to a Mg^2+^ riboswitch (Cromie et al., 2006, Dann et al., 2007) and plays a major role in the pathogenicity of *Salmonella enteric*a serovar Typhimurium (Groisman et al., 2006, Ramesh and Winkler, 2010, Groisman et al., 2013). The role of virulence for other cations such as nickel (Benoit et al., 2013), cobalt (Kersey et al., 2012, Remy et al., 2013), calcium (Sarkisova et al., 2005, Guragain et al., 2013, Sarkisova et al., 2014, Dar et al., 2016, Guragain et al., 2016, Hay et al., 2017), manganese (Boyer et al., 2002, Papp-Wallace and Maguire, 2006, Shi et al., 2014, Juttukonda and Skaar, 2015) or zinc (Dintilhac et al., 1997, Corbett et al., 2012, Mastropasqua et al., 2018, Velasco et al., 2018) has been demonstrated as well. We have not found the manganese riboswitch (*yybP-ykoY*) (Barrick et al., 2004, Dambach et al., 2015, Price et al., 2015), in spite of several known links between Mn^2+^ and virulence. Conversely, in addition to the single Ni-Co riboswitches (Furukawa et al., 2015) from the list (Hall and Lee, 2018) we have found some genes involved with nickel and/or cobalt transport associated to the cobalamin and TPP riboswitches (Table S2 and S3). Likewise, even if is not the function with which we have found it associated to a VF, ZTP-sensing has been previously associated to Zn homeostasis (Nies, 2019). Moreover, there is one hit for an Mg^2+^ ATPase C transporter found in association with the CspA thermoregulator (Table S2 and S4). These examples are indicative that genes are not always regulated by the most obvious signals, further emphasizing the importance of this compendium of VF-associated ncRNAs to provide more insight in the cues used by pathogens to regulate their virulence.

Second messengers are often involved in regulation of virulence of bacteria (Hall and Lee, 2018) and several second messengers have been shown to be sensed by riboswitches in the last decade including : cyclic-di-GMP (Sudarsan et al., 2008), cyclic-di-AMP (Nelson et al., 2013), cyclic-GAMP (Nelson et al., 2015) and ppGpp (Sherlock et al., 2018). The cyclic-di-GMP (I and II) riboswitches are known to regulate several VFs (Tamayo, 2019), but only cyclic-di-GMP-I was found in our searches. This is likely due to the relative stringency (60% identity on 98% of sequence length) when we looked for homology. Indeed, reducing our threshold to 40% identity revealed many instances of cyclic-di-GMP-II riboswitches upstream of genes encoding components of a type II secretion system, which has homologs annotated as VFs. Moreover, many other genes not recognized as VFs in PATRIC (and thus absent from our list) are also regulated by cyclic-di-GMP riboswitches, such as: *gbpA*, a characterized colonization factor from *Vibrio cholerae* (Sudarsan et al., 2008, Kariisa et al., 2016); the collagen adhesion protein from the well-known insect-killing bacteria, *Bacillus thurigiensis* (Tang et al., 2016); or several putative virulence genes from *Clostridiodes difficile* (Abt et al., 2016). Also, cyclic-di-GMP is known to influence bacterial behaviour with regards to motility or formation of biofilm, which can impact virulence (Ha and O’Toole, 2015, Valentini and Filloux, 2016), but many of the genes involved in these processes are not necessarily VFs because they are also important for the bacteria in other contexts. Thus, while we tried to be as thorough as possible, clearly the list of thousands of instances of VFs and orthologs putatively regulated by ncRNAs should not be considered as absolutely exhaustive. Other possibilities not yet annotated may also exist, e.g. while no guanidine riboswitches have been found in our search, we could presume that for bacteria which cause infection in the urinary track, guanidine riboswitches would be a good way of determining they have reached this site, and thus express relevant VFs, since guanidine is present at much higher concentration in urine (Wishart et al., 2007, Wishart et al., 2009, Duranton et al., 2012, Wishart et al., 2013, Wishart et al., 2018). Several RNA motifs known to be involved in VF regulation were not included in the present study because their annotation is deficient. Perhaps the best example for this is the RNA motif bound by the CsrA/RsmA proteins, which have a major impact on virulence (Vakulskas et al., 2015). This motif is composed of a stem-loop with a single stranded “GGA” in the loop and it is usually found in tandem where one of the two loops overlaps the ribosome binding site (RBS) (Valverde et al., 2004, Lapouge et al., 2008, Curry and Tomich, 1988, Chen et al., 1994). While our list includes the Two-AYGGAY (RF01731) family, which most likely corresponds to a subset of the 5′-UTRs targeted by CsrA/RsmA, hundreds of targets are known for these proteins (Kulkarni et al., 2014) and the binding motif consensus appears relatively relaxed, making it more difficult to annotate with a high degree of confidence. Other examples of RNA-binding protein affecting VFs exist, such as the TRAP complex which binds ∼10 repeats of (U/G)AG within one UTR (Gollnick et al., 2005), regulating genes such as *trpE* (encoding an anthranilate synthase, already shown to be regulated by TRAP (Gollnick et al., 2005), and potentially *cna3* (encoding a collagen adhesin) in *Streptococcus gallolyticus*, but which we only predicted by pattern matching (Naghdi et al., 2017) and was not confirmed experimentally as a TRAP target. We thus avoided this type of motifs for our compilation to avoid spurious annotations as much as possible.

One of the ncRNAs that was searched independently was tRNA. Many VFs on the list exhibit presence of tRNAs very close to their coding sequence (less than 30 nt). While we could not show by Northern blot that *Ef-Tu* is indeed co-transcribed with these tRNAs, they are still likely to be, given the short distance of only 46 bases separating them from the AUG. The rate of processing of the tRNAs might be too fast to permit detection of a transcript including the tRNAs together with *Ef-Tu*. In fact, co-transcription was previously observed in *E. coli* (Miyajima et al., 1981) and the proximity of tRNAs to *Ef-Tu* was already noticed in several species (Cousineau et al., 1992), which we find is generalized to numerous bacteria (Table S6), *Proteobacteria, Bacteroidetes* as well as *Firmicutes*. Presumed co-transcription of *Ef-Tu* with these tRNAs could suggest potential regulation by tRNA or merely co-regulation due to the use of the same promoter. This is further supported by the absence of predicted promoters between the tRNA closest to *Ef-Tu* and the start codon, as well as by the presence of a few promoters upstream of the three tRNA sequences, promoters which would thus also be responsible of *Ef-Tu* expression. Several roles beyond the transfer of amino acids have been demonstrated for tRNAs or fragments of tRNAs (Ryckelynck et al., 2005, Raina and Ibba, 2014, Fricker et al., 2019). The tRNA sequences found on the opposite strand, if co-transcribed with the gene downstream, could potentially be targeted by tRNAs (or tRNA fragments) in a way analogous to many sRNAs. Also, some of the machinery involved in tRNA processing and modification is known to act on mRNAs and affect their expression, like the NSun2 tRNA methylase (Zhang et al., 2012) which may imply that tRNA sequences (and pseudo-tRNAs) found in UTRs could act as substrates for such modification and processing. Furthermore, several viruses use tRNA-like motifs, either for their replication or to initiate translation in eucaryotes (Skuzeski et al., 1996, Hacker and Kaper, 2000, Zeenko et al., 2002). Finally, many bacteriophages use tRNA sequences to integrate in bacterial chromosomes, making it likely to find tRNAs in proximity to pathogenicity islands and related mobile elements (Hacker and Kaper, 2000) and implying that these tRNAs may have a critical role in horizontal gene transfer and the evolution of virulence. Yet, the role of tRNAs upstream of VFs, if any, remains to be elucidated in most cases.

Bacteria respond to signals coming from the host and its immune system. Such signal can be simple and yet present an acute change in the bacterial environment, like the change in temperature when entering a host, to which bacteria need to respond very quickly. Regulation by ncRNAs is very fast and less energetically demanding compared to regulation by protein. Discovering more ncRNAs involved in VF regulation helps better understand the means of bacteria to escape the host immune system, as well as provide potential targets to overcome bacteria pathogenicity as a promising way for treatment

## 6 Author Contributions

Bioinformatics and data analysis was performed by MRN, KS (tRNA section) and SD. Experiments were performed by MRN. Manuscript was written by MRN and JP with help from KS and SD.

## 8 Acknowledgements

We acknowledge Dr. Frédéric Veyrier (INRS-AFSB) for the kind gift of *Neisseria* RNA extracts and Emilie Boutet for her help with the Northern blot technique.

## Supplementary material for

### 1. Pipeline to noncoding RNA

1. Ribogap Query
2. Download file in CSV format/
3. Use aminoacide_extractor.pl to make Fasta format.
4. Make a blast database by indexing as explained below.
5. run Blast program “runblast_for_virulance_file.pl”
6. take the duplicate hits out by using “duplication_out.pl”.
7. Use “matrix_producer_from_excel_file.pl” to make a table with reduced result in csv format.

#### 1.1 RiboGap Query

This query should be used to find all the intergenic sequences which having non coding RNA including ribosomal RNA.

~~~
select distinct
cds.gene,cds.locus_tag,cds.product,cds.translation,cds.start as
start_of_cds,cds.end as end_of_cds,cds.strand as
strand_of_cds,fragment.fragment,fragment.description as
description_of_fragment,gap5.start as start_of_gap5,gap5.end as
end_of_gap5,gap5.strand as
strand_of_gap5,gap5.sequence,gap5.size,rna_family.fam_id,rna_family.f
am_name,rna_family.description as
description_of_rna_family,rna_family.type,rna_known.start as
start_of_rna_known,rna_known.end as end_of_rna_known,rna_known.strand
as strand_of_rna_known from fragment inner join cds on
fragment.fragment = cds.fragment inner join gap5 on cds.num_cle =
gap5.num_cle inner join rna_gap5 on gap5.num_cle = rna_gap5.num_cle
inner join rna_known on rna_gap5.rna_id = rna_known.rna_id inner join
rna_family on rna_family.fam_id = rna_known.fam_id
~~~

#### 1.2 Blast instructions

The sequences obtained from RiboGap (Naghdi et al) should be used as database. The virulence files in fasta format available from PATRIC (Wattam2014) website.

The database should be indexed by BLAST program. The following code was used to index the all Protein data from RiboGap.

~~~
instruction for indexing blast database makeblastdb -in input_db -dbtype nucl -parse_seqids
-dbtype prot
-dbtype nucl
makeblastdb -in input_db -dbtype prot -parse_seqids
~~~

#### 1.3 Scripts

~~~
**#aminoacide_extractor.pl**
#!/usr/bin/perl
use strict;
use warnings;
############################# This program extract the file downloaded from RiboGap
to take out all the AminoAcide sequences for blast
######################
############ ################################################3
use autodieqw(:all);
use DBI qw(:sql_types);
use List::AllUtilsqw(:all);
use List::Compare;
use IO::All ;
use Data::Table;
use Try::Tiny;
use File::Temp qw/ tempfiletempdir /;
use File::chmod;
use File::Copy;
################initializing the variables#########
################################# connecting to data
base#######################3333333
###################### open the file and take all the sequence from translation
sequence###################
my @file_pathogenic;
my $translation;
my @database; #### @ database stands for file dowenloded from RiboGap
my $proteine;
my $fh_out;
my $fh_error;
my $accession_locus_tag_product;
my $fasta_format;
my $count; ### to count the lines
my $product;
my $strt_cds;
my $end_cds;
my $strt_igr;
my $end_igr;
my $acc_num;
my $acc_desc;
my $locus_tag;
my $rfam_id;
my $rfam_name;
my $rfam_desc;
my $rfam_type;
my $strt_rna;
my $end_rna;
my $rna_strd;
my $gene_std;
my $gene_strd;
my $igr_srtd;
my $size_igr ;
######################################################## the path to your file location ################################3
### IN
my $path_IN=
“/home/ubuntu/Documents/RNA_disease/database_for_balst/data_25_04_2019/all_igr_ncrn a_ribogap2_25_04_2019.csv”;
#### OUT
my $path_OUT=
“/home/ubuntu/Documents/RNA_disease/database_for_balst/data_25_04_2019/translation_ aminoacide_all_igr_locus_tag_ncrna_ribogap2_24_04_2019.fasta”;
#### The path to error file
my
$path_error=“/home/ubuntu/Documents/RNA_disease/database_for_balst/data_25_04_2019/ null_protein_translation_version_24_04_2019.txt”;
###################################################################################
open ($fh_out,”>“,$path_OUT) ;
open ($fh_error,”>“, $path_error);
@file_pathogenic = io($path_IN)->slurp;
@file_pathogenic=distinct @file_pathogenic; ############# to remove duplicartion
$count=1;
foreach my $line (@file_pathogenic {
chomp $line;
unless ($count==1){ #### to omit the first line which has some text
$line =∼ s/\S+/\t/;
@database=split /\t+/, $line;
###### take the information of accession_description_locustag_product_rfam_ID_RNA_description_RNA_strand
$acc_num= $database[8];
$acc_desc =$database[9];
$locus_tag=$database[2];
$product=$database[3]; ################################## CDS ############
$strt_cds=$database[5];
$end_cds=$database[6];
$gene_strd=$database[7];
################################## IGR ###########3
$strt_igr =$database[10];
$end_igr=$database[11];
$igr_srtd= $database[12];
$size_igr =$database[14];
###################################### RFAM_RNA #########
$rfam_id =$database[15];
$rfam_name=$database[16];
$rfam_desc=$database[17];
$rfam_type=$database[18];
$strt_rna=$database[19];
$end_rna= $database[20];
$rna_strd= $database[21];
##########################################
$accession_locus_tag_product=$acc_num.”|_|”.$acc_desc.”|_|”.$locus_tag.”|_|”.$produ
ct.”|_|”.$strt_cds.”|_|”.$end_cds.”|_|”. $gene_strd.”|_|”. $strt_igr .”|_|”.$end_igr
.”|_|”. $igr_srtd .”|_|”. $rfam_id .”|_|”. $rfam_name
.”|_|”.$rfam_desc.”|_|”.$rfam_type.”|_|”. $strt_rna.”|_|”.$end_rna.”|_|” $rna_strd;
################################
$translation=$database[4];
###### check if translation is null and if so produce error file
unless ($translation eq “null”){
############################################### Print out on the monitor
print “>“, $accession_locus_tag_product, “\n”; print $translation,”\n”;
######################### print into the file ##########################
print $fh_out “>“, $accession_locus_tag_product, “\n”; print $fh_out $translation ,”\n”,
############################### the error file ########################
}else{
print $fh_error “>“, $accession_locus_tag_product, “\n”; print $fh_error $translation,”\n”,
}}
$count++;
}
close $fh_out;
close $fh_error;
print “End normal of program “,”\n”;
~~~

**#runblast_for_virulance_file.pl**

~~~
##### PATRIC
#!//usr/bin/perl
use warnings; use strict;
use List::AllUtilsqw(:all);
use Try::Tiny;
use File::Find;
use autodieqw(open system :socket);
############################## Datababes ##########################33
my
$database=“/home/ubuntu/Documents/RNA_disease/database_for_balst/data_25_04_2019/Da tabas_indexe_blast_24_04_2019/proteine_db_ribogap.fasta”;
######################################## PATRIC ##########################
my $filename=“/home/ubuntu/Documents/RNA_disease/database_for_balst/PATRIC_VF.faa”; ###########PATRIC
my $out= “/home/ubuntu/Documents/RNA_disease/database_for_balst/blast_result_version_24_04_2 019/Blast_98percent_coverage_20190614/PATRIC_result_coverege98_14_06_2019.csv”;
#################################################################################33
if (-s $filename) {
try {
## for blastp ###################### ############################## this take 98 percent coverege and not the first hit
system(“blastp -query $filename -db $database -evalue 1 -out $out -outfmt ‘6 qseqidsseqidpident length mismatch gapopenqstartqendsstart send qcovsqcovhspevaluebitscoresalltitles ‘ -qcov_hsp_perc98 -max_target_seqs 500 - threshold 11 “);
print $filename,”with “, $filename,” has finished with blast: \n”;
}catch{
  print $_,”\n”;
};
}else {
  print “sequences_query.$filename has a problem or is empty\n”;
}
print “Normal end of the script \n”;
exit;
####### VFDB
#!//usr/bin/perl
use warnings;
use strict;
use List::AllUtilsqw(:all);
use Try::Tiny;
use File::Find;
use autodieqw(open system :socket);
############################## Datababes ##########################33
my
$database=“/home/ubuntu/Documents/RNA_disease/database_for_balst/data_25_04_2019/Da
tabas_indexe_blast_24_04_2019/proteine_db_ribogap.fasta”;
################################################## VFDB #######################
my $filename=“/home/ubuntu/Documents/RNA_disease/database_for_balst/VFDB.faa”;
my $out=
“/home/ubuntu/Documents/RNA_disease/database_for_balst/blast_coverege_with_descropt ion/VFDB_coverege_description_coverege_100_07_01_2019.csv”;
if (-s $filename) {
try {
## for blastp ######################
############################## this take 98 percent coverege and not the first hit
system(“blastp -query $filename -db $database -evalue 1 -out $out -outfmt ‘6
qseqidsseqidpident length mismatch gapopenqstartqendsstart send
qcovsqcovhspevaluebitscoresalltitles ‘ -qcov_hsp_perc98 -max_target_seqs 500 - threshold 11 “);
print $filename,”with “, $filename,” has finished with blast: \n”;
}catch{
print $_,”\n”;
};
}else {
print “sequences_query.$filename has a problem or is empty\n”;
}
print “Normal end of the script \n”;
exit;
##### Victors
#!//usr/bin/perl
use warnings; use strict;
use List::AllUtilsqw(:all);
use Try::Tiny;
use File::Find;
use autodieqw(open system :socket);
############################## Datababes ##########################
my
$database=“/home/ubuntu/Documents/RNA_disease/database_for_balst/data_25_04_2019/Da
tabas_indexe_blast_24_04_2019/proteine_db_ribogap.fasta”;
######################################## Victors ###########################
my $filename=“/home/ubuntu/Documents/RNA_disease/database_for_balst/Victors.faa”;
################################ the out files could be change ########3
### Vicrtor database
my $out=
“/home/ubuntu/Documents/RNA_disease/database_for_balst/blast_result_version_24_04_2 019/Blast_98percent_coverage_20190614/Victor_result_coverege_description_coverege98
_14_06_2019.csv”;
##############################################################33
if (-s $filename) {
try {
############################# for blastp ######################
############################## this take 98 percent coverege and not the first hit
system(“blastp -query $filename -db $database -evalue 1 -out $out -outfmt ‘6 qseqidsseqidpident length mismatch gapopenqstartqendsstart send qcovsqcovhspevaluebitscoresalltitles ‘ -qcov_hsp_perc98 -max_target_seqs 500 - threshold 11 “);
print $filename,”with “, $filename,” has finished with blast: \n”;
}catch{
    print $_,”\n”;
};
}else {
print “sequences_query.$filename has a problem or is empty\n”;
}
print “Normal end of the script \n”;
exit;
~~~

**# duplication_out.pl**

~~~
#!/usr/bin/perl
use strict;
use warnings;
############################# This program take out the duplications
use autodieqw(:all);
use List::AllUtilsqw(:all);
use List::Compare;
use IO::All ;
use File::Slurp;
use Data::Table;
use Try::Tiny;
use File::Temp qw/ tempfiletempdir /;
use File::chmod;
use File::Copy;
use Text::Trim;
use Text::Table::Tiny 0.04 qw/ generate_table /;
use Text::Table::Any;
use Data::Dumper;
################initializing the variables#########
################################# connecting to data base
###open the file and take all the sequence from translation sequence##############
my @file;
my @line;
my $line;
my $fh_out;
my %seen = ();
################# referes to RF posiion on the csv file
########################################################3 the path ways
################################################### for 98 percent identity
my
$path_IN=“/home/ubuntu/Documents/RNA_disease/database_for_balst/Matrix_data/40perce
nt_id_2019_06_17”;
############### the path for output
my$path_OUT=
“/home/ubuntu/Documents/RNA_disease/database_for_balst/Matrix_data/40percent_id_201 9_06_17/without_duplication”;
##### the path for input and out put
my
$file_in=“matrix_cis_regulatory_for_all_db_40_percent_identyt_covereage_HSP_98_2019
0708.xls”;
################# 98 percent HSP and 40 percent identity CSV format ###########
my
$file_out=“uniq_matrix_cis_regulatory_for_all_db_40_percent_identyt_covereage_HSP_9
8_20190708.xls”;
####### Riboswitch 98 percent HSP and 40 percent identity CSV format ###########
open ($fh_out,”>“,”$path_OUT/$file_out”) ;
@file = io(“$path_IN/$file_in”)->tie->slurp;
#@uniq = io(“$path_OUT/$file_out”);
foreach $line (@file) {
print $fh_out $line unless $seen{$line}++;
print $line unless $seen{$line}++;
}
close $fh_out;
print “END normal”,”\n”;
~~~

**# matrix.pl**

~~~
#!/usr/bin/perl
use strict;
use warnings;
# This program extract the file downloaded from RiboGap to take out all the
AminoAcide sequences for blast
use autodieqw(:all);
use List::AllUtilsqw(:all);
use List::Compare;
use IO::All ;
use File::Slurp;
use Data::Table;
use Try::Tiny;
use File::Temp qw/ tempfiletempdir /;
use File::chmod;
use File::Copy;
use Text::Trim;
use Text::Table::Tiny 0.04 qw/ generate_table /;
use Text::Table::Any;
use Data::Dumper;
################initializing the variables#########
################################# connecting to data base
##### open the file and take all the sequence from translation sequence###########
my @file;
my @file_fp; ############# file with Fals positive RNA from RiboGap to compare
my @line;
my $fh_out;
my $fh_error;
my $fh_error_fp;
my $line_number;
################# referes to RF posiion on the csv file
my @after_RF;
my $rna_desc;
my $rfam_id;
my $rna_type;
my $accession;
my $acc_desc;
my $gene_product;
##### referr to count alle the elements number;
my %rna_id;
my %rna_type;
my %rna_descp;
my %gene_desc;
my %accession;
my %acc_num;
############################ this is just for table
my %cis_reg_table;
my %product;
my %acc_desc;
my %rfam_gene;
my %acc_count;
#######################3 cis_regulatory RNA
my %cis_reg;
my $gene_start;
my $gene_end;
my $gene_strd;
################### IGR position
my $igr_start;
my $igr_end;
my $
igr_strand;
######################### ##### RNA strand
 my $rna_strd;
 my $rna_start;
 my $rna_end;
 ############################################ TABLE ###########################
my $table;
my $row;
################################# some extra variable ##############
my %count_gene;
my $gene_num;
my %acc_desc_count;
my %tmp;
my @tmp;
########################################################## compare array ######################
my $lc;
my @intersection;
my %test;
##### This is temp path to delete later
my $path_IN=“/home/samia/Documents/virulence/new_work/result_final”;
############### to produce Matrix
##################### PATH OUT Jan 2020 ########################
##### temp path (to delete later)
my $path_OUT=“/home/samia/Documents/virulence/new_work/result_final”;
######################################## 98 percent HSP and 40 percent identity my $file_out=“matrix_riboswitch.txt”;
my
$file_out_error=“matrix_cis_regulatory_for_all_db_40_percent_identyt_covereage_HSP_
98_20190617_err.txt”;
################## Riboswitch 98 percent HSP and 40 percent identity CSV format ###########################################################################
open ($fh_out,”>“,”$path_OUT/$file_out”) ;
open ($fh_error,”>“,”$path_OUT/$file_out_error”) ;
$line_number=1;
@file = io(“$path_IN/$file_in”)->tie->chomp->slurp;
foreach my $line (@file){
###### attention to change to \t+ or “, “ ####
@line=split /\t+/, $line;
######################################## take all the element s after
RF #### take all the cis_regulatory with the same strand as gene and ncRNA
@after_RF=after_incl { $_ =∼/^RF[0-9]+/} @line;
$rfam_id=$after_RF[0];
unless ($rfam_id){
########################### The case for 224 candidate ###########3
if ($line=∼/algC/){
@after_RF=after { $_ =∼/algC/} @line;
unshift @after_RF, “RF02929” ;
$rfam_id=$after_RF[0];
$rna_desc=$after_RF[-5];
splice @after_RF, -5, 1, “$rna_desc; Cis-reg”;
$rna_desc=$after_RF[-5];
$rna_type=$after_RF[-4];
$rna_start=$after_RF[-3];
$rna_end=$after_RF[-2];
$rna_strd=$after_RF[-1];
}
elsif($line=∼/pemK/){
@after_RF=after { $_ =∼/pemK/} @line;
unshift @after_RF, “RF02913”;
$rfam_id=$after_RF[0];
$rna_desc=$after_RF[-5];
splice @after_RF, -5, 1, “$rna_desc; Cis-reg”;
$rna_desc=$after_RF[-5];
$rna_type=$after_RF[-4];
$rna_start=$after_RF[-3];
$rna_end=$after_RF[-2];
$rna_strd=$after_RF[-1];
}
elsif($line=∼/maeb/){
@after_RF=after { $_ =∼/maeb/} @line;
unshift @after_RF, “RF0maeb” ;
$rfam_id=$after_RF[0];
$rna_desc=$after_RF[-5];
splice @after_RF, -5, 1, “$rna_desc; Cis-reg”;
$rna_desc=$after_RF[-5];
$rna_type=$after_RF[-4];
$rna_start=$after_RF[-3];
$rna_end=$after_RF[-2];
$rna_strd=$after_RF[-1];
}
Elsif($line=∼/DUF1646/){
@after_RF=after { $_ =∼/DUF1646/} @line;
unshift @after_RF, “RF03071” ;
$rfam_id=$after_RF[0];
$rna_desc=$after_RF[-5];
splice @after_RF, -5, 1, “$rna_desc; Cis-reg”;
$rna_desc=$after_RF[-5];
$rna_type=$after_RF[-4];
$rna_start=$after_RF[-3];
$rna_end=$after_RF[-2];
$rna_strd=$after_RF[-1];
}
Elsif($line=∼/malK-I/){
@after_RF=after { $_ =∼/malK-I/} @line;
unshift @after_RF, “RF03069” ;
$rfam_id=$after_RF[0];
$rna_desc=$after_RF[-5];
splice @after_RF, -5, 1, “$rna_desc; Cis-reg”;
$rna_desc=$after_RF[-5];
$rna_type=$after_RF[-4];
$rna_start=$after_RF[-3];
$rna_end=$after_RF[-2];
$rna_strd=$after_RF[-1];
}
Elsif($line=∼/narK/){
@after_RF=after{ $_ =∼/narK/} @line;
unshift @after_RF, “RF03032” ;
$rfam_id=$after_RF[0];
$rfam_id=$after_RF[0];
$rna_desc=$after_RF[-5];
splice @after_RF, -5, 1, “$rna_desc; Cis-reg”;
$rna_desc=$after_RF[-5];
$rna_type=$after_RF[-4];
$rna_start=$after_RF[-3];
$rna_end=$after_RF[-2];
$rna_strd=$after_RF[-1];
}
elsif($line=∼/Rothia-sucC/){
@after_RF=after { $_ =∼/Rothia-sucC/} @line; unshift @after_RF, “RF03024” ;
$rfam_id=$after_RF[0];
$rna_desc=$after_RF[-5];
splice @after_RF, -5, 1, “$rna_desc; Cis-reg”;
$rna_desc=$after_RF[-5];
$rna_type=$after_RF[-4];
$rna_start=$after_RF[-3];
$rna_end=$after_RF[-2];
$rna_strd=$after_RF[-1];
}
elsif($line=∼/sul1/){
@after_RF=after { $_ =∼/sul1/} @line;
unshift @after_RF, “RF03058” ;
$rfam_id=$after_RF[0];
$rna_desc=$after_RF[-5];
splice @after_RF, -5, 1, “$rna_desc; Cis-reg”;
$rna_desc=$after_RF[-5];
$rna_type=$after_RF[-4];
$rna_start=$after_RF[-3];
$rna_end=$after_RF[-2];
$rna_strd=$after_RF[-1];
}
elsif($line=∼/terC/){
@after_RF=after { $_ =∼/terC/} @line;
unshift @after_RF, “RF03067” ;
$rfam_id=$after_RF[0];
$rna_desc=$after_RF[-5];
splice @after_RF, -5, 1, “$rna_desc; Cis-reg”;
$rna_desc=$after_RF[-5];
$rna_type=$after_RF[-4];
$rna_start=$after_RF[-3];
$rna_end=$after_RF[-2];
$rna_strd=$after_RF[-1];
}
else{
print $fh_error “This line has problem of RFAM id :\n”, join “\t”, @line, “\n”;
next;
 }
}else{
$rna_desc=$after_RF[12];
$rna_type=$after_RF[13];
$rna_start=$after_RF[14];
$rna_end=$after_RF[15];
$rna_strd=$after_RF[16];
$accession= $after_RF[1];
$acc_desc= $after_RF[2];
$gene_product=$after_RF[4];
$gene_start=$after_RF[5];
$gene_end=$after_RF[6];
$gene_strd=$after_RF[7]; #################### IGR position
$igr_start=$after_RF[8] ;
$igr_end=$after_RF[9];
$igr_strand=$after_RF[10];
}
#################### attention for RNA_decs and RNA_ type ######################## ######################### RNA type is for all RFAMid
## RNA desc is for all nonRFAMid motif (224 motif) at the time of writing this code
#### you can choose for what you are looking for
###for 224 motif you should choose rna _typ instead of rna_desc (RNA description)
# if ($rna_desc=∼/Cis-reg[A-Za-z]?/ and $gene_strd == $rna_strd){ # if ($rna_desc=∼/riboswitch/ and $gene_strd == $rna_strd){
if ($rna_type=∼/riboswitch/ and $gene_strd == $rna_strd){
# if ($rna_type=∼/Cis-reg[A-Za-z]?/ and $gene_strd == $rna_strd){ # if ($rna_desc=∼/(thermo.*)/ and $gene_strd == $rna_strd){
#if ($rna_desc=∼/(TPP riboswitch)/ and $gene_strd == $rna_strd){ #if ($rna_desc=∼/(NiCo riboswitch)/ and $gene_strd == $rna_strd){
#if ($rna_desc=∼/(NiCo riboswitch)|(Glycine riboswitch)/ and $gene_strd ==
$rna_strd){
#if ($rna_desc=∼/(TPP riboswitch)|(FMN riboswitch)|(Glycine riboswitch)/ and
$gene_strd == $rna_strd){
#if ($rna_desc=∼/(TPP riboswitch)|(NiCo riboswitch)|(FMN riboswitch)/ and
$gene_strd == $rna_strd){
#if ($rna_desc=∼/(TPP riboswitch)|(SAM riboswitch)/ and $gene_strd == $rna_strd
){
################ only riboswitch #######################
$rna_id{$rfam_id}++;
push (@{$product{$rfam_id}},$gene_product); push (@{$accession{$rfam_id}},$acc_desc); push (@{$acc_num{$rfam_id}},$accession);
push (@{$acc_desc{$rfam_id}},$accession.”_”.$acc_desc); push (@{$cis_reg{$rfam_id}},$accession.”_”.$acc_desc); push (@{$cis_reg_table{$rfam_id}},$rna_desc, $rna_type);
##### this is for having distinct Rfam id @{$cis_reg{$rfam_id}}=distinct (@{$cis_reg{$rfam_id}});
##### this is for having the distict accession descriptipn @{$accession{$rfam_id}}=distinct (@{$accession{$rfam_id}});
##### this is for having the distictaccession_description number @{$acc_desc{$rfam_id}}=distinct(@{$acc_desc{$rfam_id}});
##### this is for having the distictrna_desc_for table @{$cis_reg_table{$rfam_id}}=distinct(@{$cis_reg_table{$rfam_id}});
##### this is for having the distictaccesion_number
}
$line_number++;
}
########################### cis_regulatory ##################################3
##### first count the gene number then do your table #foreach my $k (sort keys %cis_reg){
foreach my $k (sort keys %acc_desc){ my $gene_num;
###################### this is for the table
foreach $gene_product(distinct (@{$product{$k}})){ foreach $gene_num (@{$product{$k}}){
$count_gene{$gene_num}++;
}
push (@$row ,[ $k,(join “;”, @{$cis_reg_table{$k}}), $rna_id{$k}, $gene_product
,scalar @{$product{$k}}, $count_gene {$gene_product}, (shift @{$acc_desc{$k}})]);
%count_gene=();
}
}
###################### add the header for table ######################3
#unshift (@$row, [ “Rfam_id”,”RNA_description”, “RNA_instance “, “gene_product”, “Gene_instance”, “Accession_desc”]);
#unshift (@$row, [ “Rfam_id”,”RNA_description”, “RNA_instance “, “gene_product”, “Gene_instance”, “Accession”,”Accession_desc”, “Accession_count”]);
#unshift (@$row, [ “Rfam_id”,”RNA_description”, “RNA_instance “,”gene_rna”, “gene_product”, “Gene_instance”]);
#unshift (@$row, [ “Rfam_id”,”RNA_description”, “RNA_instance “, “test”, “gene_product”, “Gene_instance”]);
#unshift (@$row, [ “Rfam_id”,”RNA_description”, “RNA_instance (total) “, “gene_product”, “Gene_instance”]);
#unshift (@$row, [ “Rfam_id”,”RNA_description”, “RNA_instance (total) “, “gene_product”, “Gene_instance(total)”, “Accession_description”]);
#unshift (@$row, [ “Rfam_id”,”RNA_description”, “RNA_instance (total) “, “gene_product”, “Associate_gene_for_this_rfam_id”, “total_found_gene”, “Accession_description”]);
unshift (@$row, [ “Rfam_id”,”RNA_description”, “RNA_instance (total) “, “gene_product”, “Associate_gene_for_this_rfam_id”, “total_found_gene”, “Accession_description”]);
push @$table, @$row;
######################## Print in TSV format ##################################
print Text::Table::Any::table(rows => $table, header_row => 1,
backend => ‘Text::Table::TSV’);
print $fh_outText::Table::Any::table(rows => $table, header_row => 1,
backend => ‘Text::Table::TSV’);
close $fh_out;
close $fh_error;
#close $
fh_error_fp;
print “END normal of Program”,”\n”;
~~~

### Evaluating co-transcription of tRNAs with Ef-Tu

#### Sequences of Neisseria species evaluated

~~~
**>Neisseria_gonorrhoeae_F62_cont1_29_whole genome shotgun sequence**
ATGGCTAAGGAAAAATTCGAACGTAGCAAACCGCACGTAAACGTTGGCACCATCGGTCACGTTGACCATGGTAAAACCACCCTGACTGCCGCTTTGACTACTA TTTTAGCTAAAAAATTCGGCGGCGCTGCAAAAGCTTACGACCAAATCGACAACGCACCCGAAGAAAAAGCACGCGGTATTACCATTAACACCTCGCACGTAGA ATACGAAACCGAAACCCGCCACTACGCACACGTAGACTGTCCGGGTCACGCCGACTACGTTAAAAACATGATTACCGGCGCCGCACAAATGGACGGTGCAATC CTGGTATGTTCTGCTGCCGACGGCCCTATGCCGCAAACCCGCGAACACATCCTGCTGGCCCGTCAAGTAGGCGTACCTTACATCATCGTGTTCATGAACAAAT GCGACATGGTCGACGATGCCGAGCTGTTGGAACTGGTTGAAATGGAAATCCGCGACCTGCTGTCCAGCTACGACTTCCCCGGCGACGACTGCCCGATCGTACA AGGTTCCGCACTGAAAGCCTTGGAAGGCGATGCCGCTTACGAAGAAAAAATCTTCGAACTGGCTACCGCATTGGACAGCTACATCCCGACTCCCGAGCGTGCC GTGGACAAACCATTCCTGCTGCCTATCGAAGACGTGTTCTCCATTTCCGGCCGCGGTACCGTAGTCACCGGCCGTGTAGAGCGAGGTATCATCCACGTTGGTG ACGAGATTGAAATCGTCGGTCTGAAAGAAACCCAAAAAACCACCTGTACCGGCGTTGAAATGTTCCGCAAACTGCTGGACGAAGGTCAGGCGGGCGACAACGT AGGCGTATTGCTGCGCGGTACCAAACGTGAAGACGTAGAACGCGGTCAGGTATTGGCCAAACCGGGTACTATCACTCCTCACACCAAGTTCAAAGCAGAAGTG TACGTATTGAGCAAAGAAGAGGGCGGCCGCCATACCCCGTTTTTCGCCAACTACCGTCCCCAATTCTACTTCCGTACCACTGACGTAACCGGCGCGGTTACTT TGGAAAAAGGTGTGGAAATGGTAATGCCGGGTGAGAACGTAACCATTACTGTAGAACTGATTGCGCCTATCGCTATGGAAGAAGGTCTGCGCTTTGCGATTCG CGAAGGCGGCCGTACCGTGGGTGCCGGCGTGGTTTCTTCTGTTATCGCTTAA
**>NC_003112.2_Neisseria meningitidis MC58_NMB0124_elongation factor Tu**
ATGGCTAAGGAAAAATTCGAACGTAGCAAACCGCACGTAAACGTTGGCACCATCGGTCACGTTGACCATGGTAAAACCACCCTGACTGCCGCTTTGACTACTA TTTTGGCTAAAAAATTCGGCGGTGCTGCAAAAGCTTACGACCAAATCGACAACGCACCCGAAGAAAAAGCACGCGGTATTACCATTAACACCTCGCACGTGGA ATACGAAACCGAAACCCGCCACTACGCACACGTAGACTGCCCGGGGCACGCCGACTACGTTAAAAACATGATTACCGGCGCCGCACAAATGGACGGTGCAATC CTGGTATGTTCCGCAGCCGACGGCCCTATGCCGCAAACCCGCGAACACATCCTGCTGGCCCGCCAAGTAGGCGTACCTTACATCATCGTGTTCATGAACAAAT GCGACATGGTCGACGATGCCGAGCTGTTGGAACTGGTTGAAATGGAAATCCGCGACCTGCTGTCCAGCTACGACTTCCCCGGCGATGACTGCCCGATTGTACA AGGTTCCGCACTGAAAGCCTTGGAAGGCGATGCCGCTTACGAAGAAAAAATCTTCGAACTGGCTGCCGCATTGGACAGCTACATCCCGACTCCCGAGCGAGCC GTGGACAAACCGTTCCTGCTGCCTATCGAAGACGTGTTCTCCATTTCCGGCCGCGGTACAGTAGTAACCGGCCGTGTAGAGCGCGGTATCATCCACGTTGGTG ACGAGATTGAAATCGTCGGTCTGAAAGAAACCCAAAAAACCACTTGTACCGGTGTTGAAATGTTCCGCAAACTGCTGGACGAAGGTCAGGCGGGCGACAACGT AGGCGTATTGCTGCGCGGTACCAAACGTGAAGACGTGGAACGCGGTCAGGTATTGGCTAAACCGGGTACTATCACTCCTCACACCAAATTCAAAGCAGAAGTA TACGTACTGAGCAAAGAAGAGGGTGGTCGTCACACTCCGTTCTTCGCCAACTACCGTCCGCAATTCTACTTCCGTACCACCGACGTAACCGGCGCGGTTACTT TGGAAGAAGGTGTAGAAATGGTAATGCCGGGTGAAAACGTAACCATCACCGTAGAACTGATTGCGCCTATCGCTATGGAAGAAGGCCTGCGCTTTGCGATTCG CGAAGGCGGCCGTACCGTGGGTGCCGGCGTGGTTTCTTCTGTTATCGCTTAA
**>NZ_CP020452.2_Neisseriasicca_locus_tag_A6J88_RS18225_Eftu**
ATGGCTAAGGAAAAATTTGAACGTAGCAAACCGCACGTAAACGTTGGCACCATCGGTCACGTTGACCATGGTAAAACCACCCTAACTGCTGCTTTGACTACTA TTTTGGCTAAAAAATTCGGCGGTGCTGCAAAAGCTTACGACCAAATCGACAACGCTCCCGAAGAAAAAGCCCGCGGTATTACCATCAATACCTCACACGTAGA ATACGAAACCGAAACCCGCCACTACGCACACGTAGACTGCCCGGGTCACGCCGACTACGTTAAAAACATGATTACCGGTGCCGCACAAATGGACGGCGCAATC TTGGTATGTTCTGCTGCCGACGGTCCTATGCCGCAAACCCGCGAACACATCCTGTTGGCCCGTCAAGTAGGTGTACCTTACATCATCGTGTTCATGAACAAAT GCGACATGGTTGACGATGCCGAATTGTTGGAACTGGTTGAAATGGAAATCCGTGACTTGCTGTCAAGCTACGACTTCCCTGGTGACGACTGCCCGATTGTACA AGGTTCTGCACTGAAAGCCTTGGAAGGCGATGCCGCTTACGAAGAAAAAATCTTCGAACTGGCTGCCGCATTGGACAGCTACATCCCGACTCCCGAGCGTGCC GTAGACAAACCGTTCCTGTTGCCTATCGAAGACGTATTCTCCATCTCCGGTCGTGGTACAGTAGTAACCGGCCGTGTAGAGCGCGGTGTTATCCACGTTGGTG ACGAGATCGAAATCGTAGGTCTGAAAGAAACCCAAAAAACCACTTGTACCGGTGTTGAAATGTTCCGCAAACTGCTGGACGAAGGTCAAGCAGGTGACAACGT AGGCGTATTGCTGCGCGGTACCAAACGTGAAGAAGTGGAACGCGGTCAAGTATTGGCTAAACCGGGTACCATCACTCCGCACACCAAATTCAAAGCAGAAGTG TACGTATTGAGCAAAGAAGAGGGTGGTCGTCATACTCCGTTCTTCGCTAACTACCGTCCTCAATTCTACTTCCGTACTACCGACGTAACCGGCGCGGTTACTT TGGAAGAAGGTGTAGAAATGGTTATGCCTGGTGAGAACGTAGCCATCACTGTAGAATTGATTGCACCTATCGCTATGGAAGAAGGTCTGCGCTTTGCGATTCG CGAAGGTGGCCGTACCGTAGGTGCAGGCGTGGTTTCTTCTATCATCGCTTAA
**>NZ_CP007726.1_Neisseriaelongata subsp. glycolytica ATCC _29315_locus_tag_NELON_RS0103_elongation factor Tu**
ATGGCAAAGGAAAAATTTGAACGTAGCAAACCGCACGTAAACGTTGGCACCATCGGTCACGTTGACCATGGTAAAACCACTCTGACTGCTGCTTTGACTACTA TTTTGGCTAAAAAATTCGGTGGTGCTGCAAAAGCCTACGATCAAATCGACAACGCTCCCGAAGAAAAAGCCCGCGGTATTACCATTAATACCTCACACGTAGA ATACGAAACCGAAACCCGCCACTACGCACACGTAGACTGCCCGGGTCACGCCGACTACGTTAAAAATATGATTACCGGTGCCGCACAAATGGACGGCGCAATC TTGGTATGTTCCGCTGCTGACGGTCCTATGCCGCAAACCCGCGAACACATCCTGTTGGCCCGCCAAGTAGGCGTACCTTACATCATCGTGTTTATGAACAAAT GCGACATGGTTGACGATGCCGAGCTGTTGGAACTGGTTGAAATGGAAATCCGTGACTTGCTGTCAAGCTACGACTTCCCAGGTGATGACTGCCCGATCGTACA AGGTTCCGCACTGAAAGCCTTGGAAGGCGACGCAGCTTACGAAGAAAAAATCTTCGAACTGGCTGCCGCATTGGATAGCTACATCCCGACTCCTGAGCGTGCC GTGGACAAACCGTTCCTGTTGCCTATCGAAGACGTATTCTCCATCTCCGGTCGTGGTACAGTAGTAACCGGTCGTGTAGAGCGTGGTGTTATCCACGTTGGTG ACGAGATCGAAATCGTAGGTCTGAAAGAAACCCAAAAAACCACTTGTACCGGTGTTGAGATGTTCCGCAAACTGCTGGACGAGGGTCAAGCCGGTGACAACGT AGGCGTATTGCTGCGCGGTACCAAACGCGAAGAAGTGGAACGCGGTCAAGTATTGGCTAAACCGGGTACCATCACTCCGCACACCAAATTCAAAGCAGAAGTG TACGTGTTGAGCAAAGAAGAGGGTGGTCGTCATACTCCGTTCTTCGCTAACTACCGTCCACAATTCTACTTCCGTACTACCGACGTAACCGGCGCGGTTACTT TGGAAGAAGGTGTAGAAATGGTTATGCCTGGTGAGAACGTAGCCATCACTGTAGAACTGATTGCACCGATCGCTATGGAAGAAGGTCTGCGCTTTGCGATTCG CGAAGGTGGCCGTACCGTAGGTGCGGGCGTGGTTTCTTCTGTTATCGCTTAA
~~~

**Table S1:**
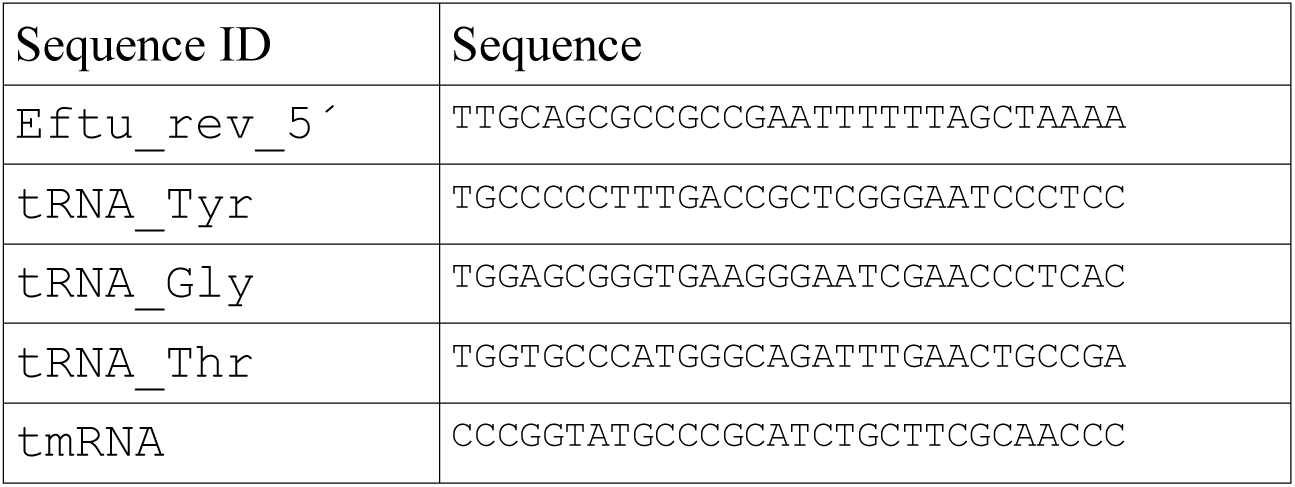
The primer sequences used for Northern blot

**Figure S1.**
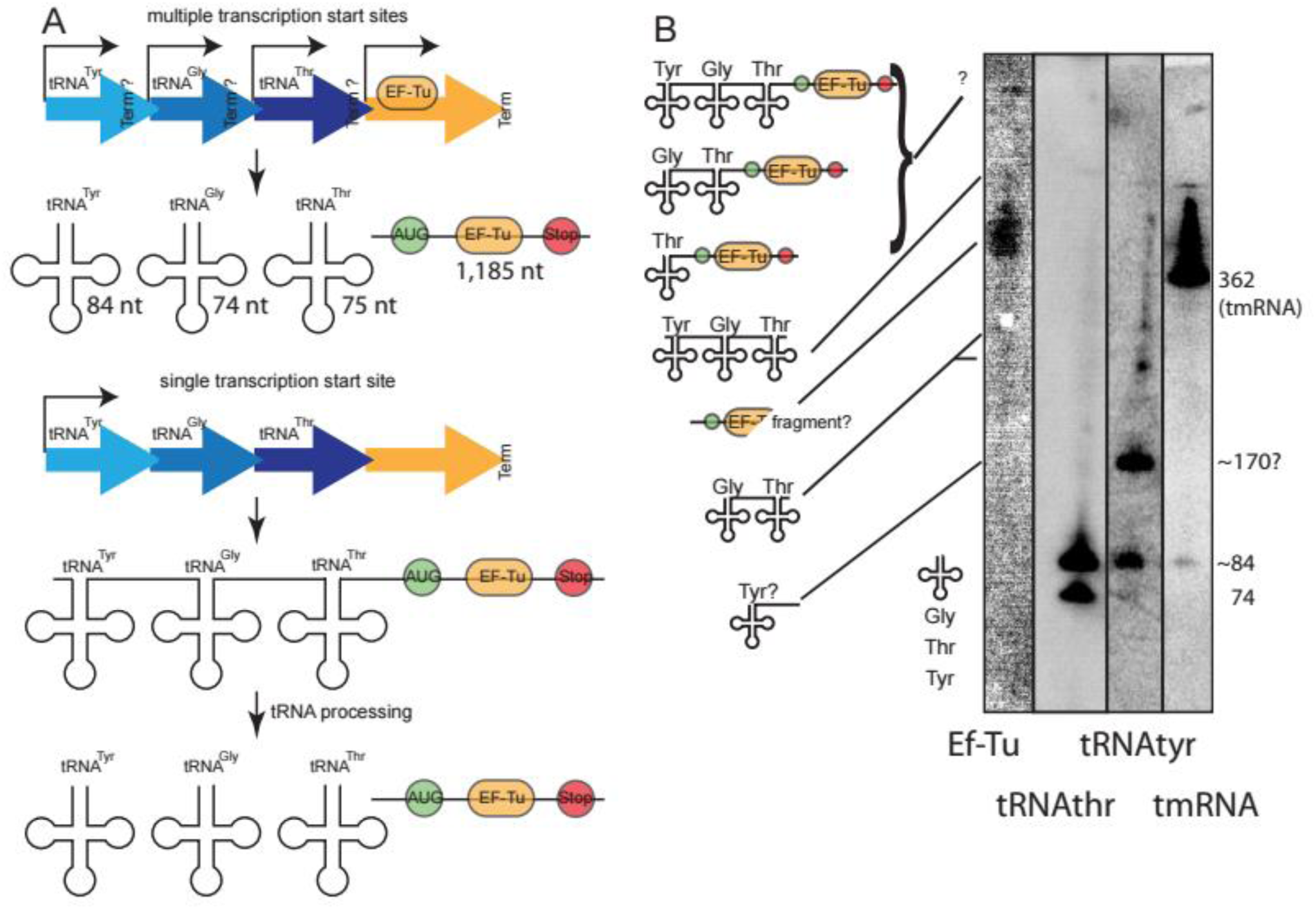
Co-transcription of tRNAs upstream of Ef-Tu. **A)** Potential outcomes of transcription. **B)** Co-transcription of Ef-Tu and tRNAs upstream was evaluated by northern blot.Total RNA from *Neisseria gonorrhoeae* was used and probed with sequences in table S1 for the RNAs as indicated under the Northern image. Detection of transcripts with, presumably, multiple tRNAs could be performed, but not with Ef-Tu. This is probably due to the rapid processing of tRNAs.The tmRNA was used as a size control (it should be 362 bases).

**Figure S2.**
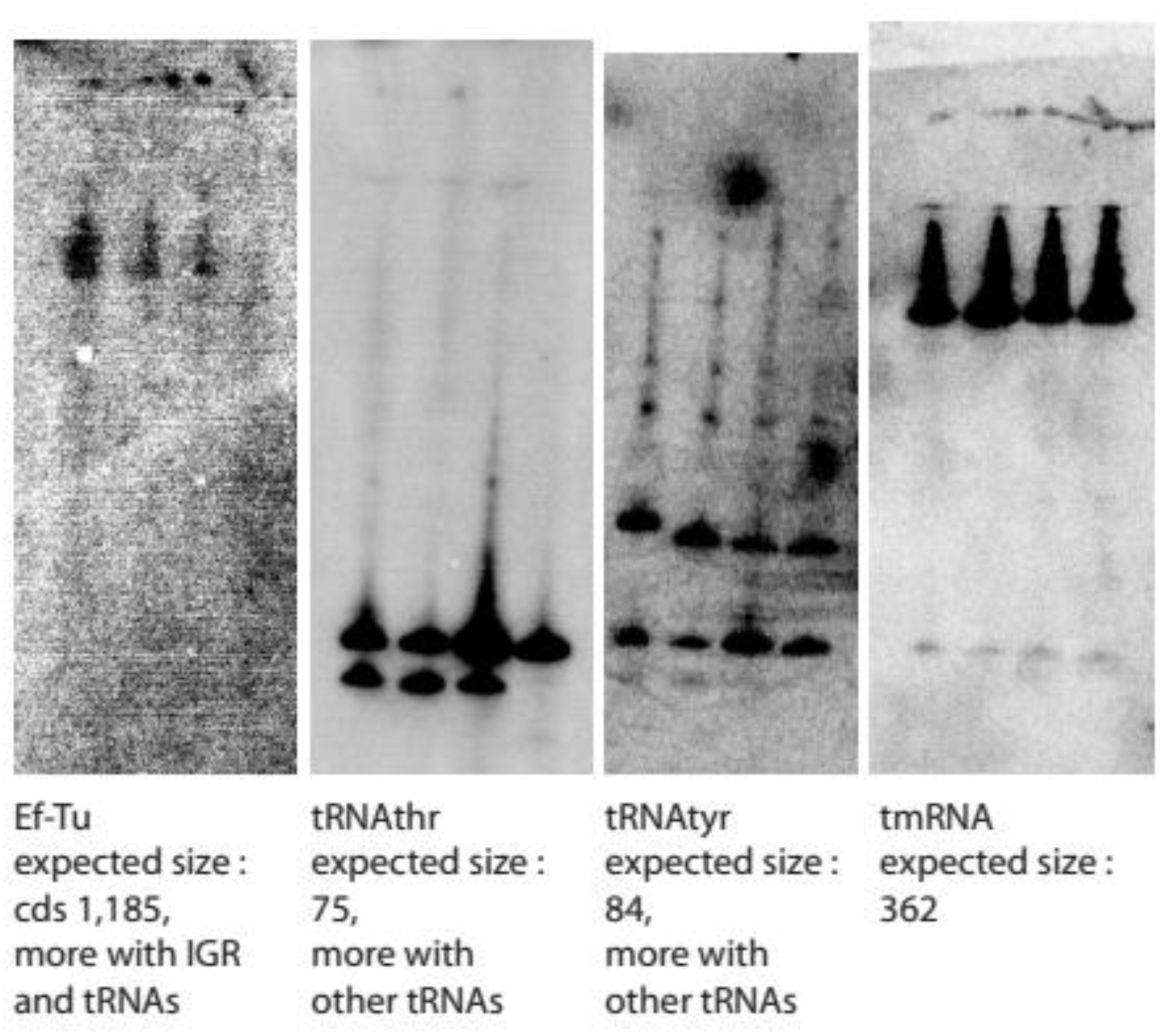
Full Northerns of figure S1. Probed were used in the same order presented in figure. Wells correspond to total RNA from: *Neisseria gonorrhoeae, Neisseria meningitidis, Neisseria sicca and Neisseria elongata*, in this order.This is a representative membrane (out of three experiments).

In summary, while very weak bands corresponding to transcripts not fully processed with the tRNA^thr^ and tRNA^tyr^ probes, no convincing band could be detected that would have had the expected size of the Ef-Tu mRNA + one or more tRNAs still part of the same transcript. We have also tried amplifying such transcripts by RT-PCR without success. These results indicate that if Ef-Tu is co-transcribed with tRNAs (as suggested by this genetic arrangement), the transcript is quickly processed in its individual genes (as it is also suggested by the overwhelming intensity of bands corresponding to processed tRNA compared to the bands putatively comprising two or three tRNAs). Nevertheless, this does not preclude that Ef-Tu may use one of the tRNAs’ promoter(s) for its transcription.

(Tables S2, S3, S4, S5 and S6 are in separate excel files.)

### tRNAs upstream of VFs (and homologs)

(Table S6 is in a separate excel file.)

Additional methodology details

- Profile of tRNAs upstream of virulence genes: We evaluated the distribution of tRNAs upstream of the virulence genes, which were compared to 100 genes randomly taken from the genome of the strain Escherichia coli K12 MG1655. The 100 genes were randomly selected with Excel’s RAND () function in three replicas For each replica, the 5’ UnTranslated Reagion (5’UTR) were extracted from the Ribogap database (http://ribogap.iaf.inrs.ca) following mysql requests (see supplementary material, previous section).
- Method
  - 5’UTRs sequences: extraction from the ribogap database (see the mysql scripts in the additional material)
  - Analysis with tRNAscan-SE (Chan and Lowe, 2019) (http://lowelab.ucsc.edu/tRNAscan-SE/): tRNAscan is a tool for detecting tRNA genes and predicting function from genomic sequences. We used the locally installed source code version to make our predictions. The configuration parameters were left by default (see material additional).

**Table S7:** Virulence genes with pseudo-tRNA predicted in their 5’UTR

| Gene | Nbr pseudo-tRNA | nbr 5'UTR |
| --- | --- | --- |
| acpP1 | 1 | 4650 |
| aroQ | 1 | 2499 |
| clpP | 14 | 412 |
| cysM | 1 | 2218 |
| gtrA | 3 | 143 |
| mcp2 | 2 | 74 |
| nadA | 5 | 2187 |
| pfoR | 1 | 88 |
| pgmB | 1 | 613 |
| pyrE | 1 | 4989 |
| secE | 1 | 8284 |
| tig | 1 | 6718 |
| tuf | 1 | 9187 |
| tufB | 1 | 596 |
| vacB | 1 | 625 |

**Table S8:** Overall tRNAscan results

|  | Random genes |  |  |  |  | Virulence genes |
| --- | --- | --- | --- | --- | --- | --- |
|  | Replica 1 | Replica 2 | Replica 3 | Mean | Standard deviation |  |
| *5'UTRs* | 69456 | 86744 | 84256 | 80152 | 9346.17 | 1345678 |
| *5'UTRs with tRNA* | 2074 | 2124 | 2375 | 2191 | 161.3 | 3971 |
| *5'UTRs with pseudo tRNA* | 6 | 36 | 54 | 32 | 24.25 | 35 |
| *genes* | 100 | 100 | 100 | 100 | 0 | 7042 |
| *genes with tRNA* | 57 | 60 | 47 | 55 | 6.8 | 426 |
| *genes with pseudo tRNA* | 1 | 4 | 7 | 4 | 3 | 15 |

**Figure S3.**
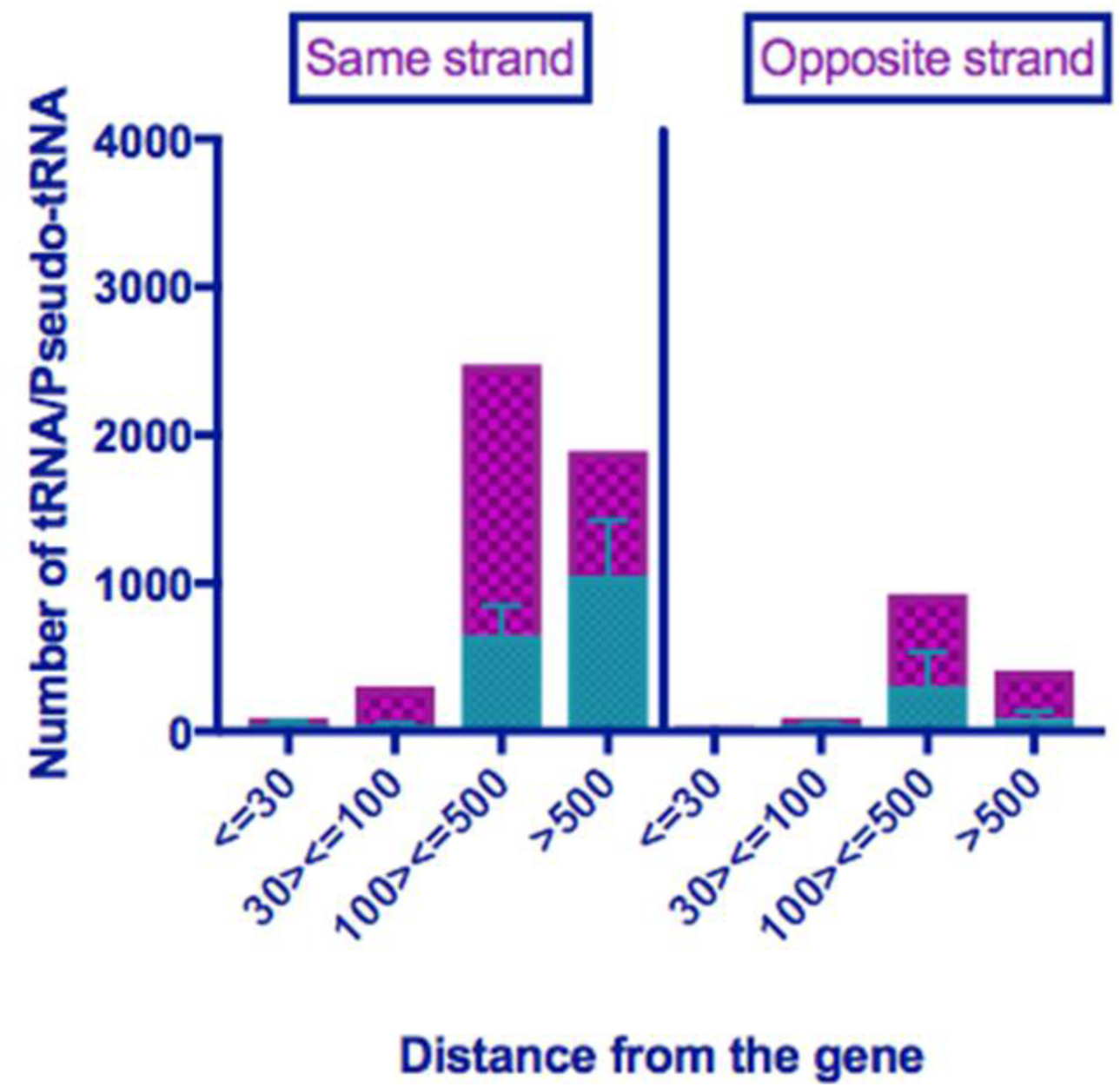
Distribution and orientation of tRNA (and pseudo tRNA) sequences upstream of genes. Number of tRNAs upstream of VFs (or homologs) in purple, number of tRNAs upstream of randomized controls in turquoise.

While there are numerous tRNAs found upstream of VFs (and homologs), they are not more abundant, proportionally, to the randomized controls. Nevertheless, no particular general trend was expected and the significance of these genetic arrangements, especially for those that are very close to the coding sequence, merits inspection on a case by case basis.

### Transcription terminators

(Tables S9 and S10 are in separate excel files.)

**Table S11:** The genes which have rho independent terminator

| **locus_tag** | **Gene** | **Start** | **end** |
| --- | --- | --- | --- |
| BSU04490 | putative ABC transporter | 502668 | 502883 |
| BSU06360 | GMP synthetase | 692609 | 692701 |
| BSU07260 | lipoteichoic acid synthase | 796130 | 796224 |
| BSU09590 | putative membrane protein | 1035409 | 1035538 |
| BSU18090 | subunit B of DNA topoisomerase IV | 1933177 | 1933398 |
| lmo0517 | Phosphoglycerate mutase | 552417 | 552315 |
| lmo2187 | Protein of unknown function | 2275362 | 2275261 |
| EF1147 | CTP synthetase | 1115883 | 1115958 |

These locus_tag have Rho-independent transcription terminators RiTTs just upstream of VFs (or VF homologs) as indicated by our search. They also correspond to experimentally determined premature transcription termination sites in *Listeria monocytogenes* as determined by Term-seq (Dar et al., 2016). This highlights the usefulness of using the RiTTs and RTTs to quickly survey potential transcription-mediated mechanisms of gene regulation.

## References

Abduljalil, J. (2018). Bacterial riboswitches and RNA thermometers: Nature and contributions to pathogenesis. Noncoding RNA Res. 3, 54–63.

Abt, M., McKenney, P. and Pamer, E. (2016). Clostridium difficile colitis: pathogenesis and host defence. Nat. Rev. Microbiol. 14, 609–620.

Altschul, S., Gish, W., Miller, W., Myers, E. and Lipman, D. (1990). Basic local alignment search tool. J. Mol. Biol. 215, 403–410.

Barrick, J., Corbino, K., Winkler, W., Nahvi, A., Mandal, M., Collins, J. et al. (2004). New RNA motifs suggest an expanded scope for riboswitches in bacterial genetic control. Proc. Natl. Acad. Sci. U.S.A. 101, 6421–6426.

Bäumler, A. and Sperandio, V. (2016). Interactions between the microbiota and pathogenic bacteria in the gut. Nature 535, 85-93. doi: 10.1038/nature18849.

Benoit, S., Miller, E. and Maier, R. (2013). Helicobacter pylori stores nickel to aid its host colonization. Infect. Immun. 81, 580–584.

Boyer, E., Bergevin, I., Malo, D., Gros, P. and Cellier, M. (2002). Acquisition of Mn(II) in addition to Fe(II) is required for full virulence of Salmonella enterica serovar Typhimurium. Infect. Immun. 70, 6032–6042.

Broder, U., Jaeger, T. and Jenal, U. (2016). LadS is a calcium-responsive kinase that induces acute-to-chronic virulence switch in Pseudomonas aeruginosa. Nat Microbiol. 2, 16184.

Brown, S., Cornforth, D. and Mideo, N. (2012). Evolution of virulence in opportunistic pathogens: generalism, plasticity, and control.. Trends Microbiol. 20, 336-342. doi: 10.1016/j.tim.2012.04.005.

Caldelari, I., Chao, Y., Romby, P. and Vogel, J. (2013). RNA-mediated regulation in pathogenic bacteria.. Cold Spring Harb Perspect Med 3, a010298. doi: 10.1101/cshperspect.a010298.

Camacho, C., Coulouris, G., Avagyan, V., Ma, N., Papadopoulos, J., Bealer, K. et al. (2009). BLAST+: architecture and applications. BMC Bioinformatics 10, 421.

Casadevall, A. and Pirofski, L. (2003). The damage-response framework of microbial pathogenesis.. Nat Rev Microbiol. 1, 17-24. doi: 10.1038/nrmicro732.

Chakravarty, S. and Massé, E. (2019). RNA-Dependent Regulation of Virulence in Pathogenic Bacteria. Front Cell Infect Microbiol. 9, 337.

Chan, P. and Lowe, T. (2019). tRNAscan-SE: Searching for tRNA Genes in Genomic Sequences. Methods Mol. Biol. 1962, 1–14.

Chen, H., Bjerknes, M., Kumar, R. and Jay, E. (1994). Determination of the optimal aligned spacing between the Shine-Dalgarno sequence and the translation initiation codon of Escherichia coli mRNAs. Nucleic Acids Res. 22, 4953–4957.

Chen, L., Zheng, D., Liu, B., Yang, J. and Jin, Q. (2016). VFDB 2016: hierarchical and refined dataset for big data analysis--10 years on.. Nucleic Acids Res. 44, D694-D697. doi: 10.1093/nar/gkv1239.

Clarke, M., Hughes, D., Zhu, C., Boedeker, E. and Sperandio, V. (2006). The QseC sensor kinase: a bacterial adrenergic receptor. Proc. Natl. Acad. Sci. U.S.A. 103, 10420–10425.

Corbett, D., Wang, J., Schuler, S., Lopez-Castejon, G., Glenn, S., Brough, D. et al. (2012). Two zinc uptake systems contribute to the full virulence of Listeria monocytogenes during growth in vitro and in vivo. Infect. Immun. 80, 14–21.

Cousineau, B., Cerpa, C., Lefebvre, J. and Cedergren, R. (1992). The sequence of the gene encoding elongation factor Tu from Chlamydia trachomatis compared with those of other organisms. Gene 120, 33–41.

Cromie, M., Shi, Y., Latifi, T. and Groisman, E. (2006). An RNA sensor for intracellular Mg(2+). Cell 125, 71–84.

Curry, K. and Tomich, C. (1988). Effect of ribosome binding site on gene expression in Escherichia coli. DNA 7, 173–179.

Dambach, M., Sandoval, M., Updegrove, T., Anantharaman, V., Aravind, L., Waters, L. et al. (2015). The ubiquitous yybP-ykoY riboswitch is a manganese-responsive regulatory element. Mol. Cell 57, 1099–1109.

Dann, C., Wakeman, C., Sieling, C., Baker, S., Irnov, I. and Winkler, W. (2007). Structure and mechanism of a metal-sensing regulatory RNA. Cell 130, 878–892.

Dar, D., Shamir, M., Mellin, J., Koutero, M., Stern-Ginossar, N., Cossart, P. et al. (2016). Term-seq reveals abundant ribo-regulation of antibiotics resistance in bacteria. Science 352, aad9822.

Dintilhac, A., Alloing, G., Granadel, C. and Claverys, J. (1997). Competence and virulence of Streptococcus pneumoniae: Adc and PsaA mutants exhibit a requirement for Zn and Mn resulting from inactivation of putative ABC metal permeases. Mol. Microbiol. 25, 727–739.

DiRita, V., Engleberg, N., Heath, A., Miller, A., Crawford, J. and Yu, R. (2000). Virulence gene regulation inside and outside. Philos Trans R Soc Lond B Biol Sci. 355, 657-665. doi: 10.1098/rstb.2000.0606.

Di Salvo, M., Puccio, S., Peano, C., Lacour, S., Alifanoe, P. (2019) RhoTermPredict: an algorithm for predicting Rho-dependent transcription terminators based on Escherichia coli, Bacillus subtilis and Salmonella enterica databases. BMC Bioinformatics 20, 117.

Do, H., Makthal, N., VanderWal, A., Saavedra, M., Olsen, R., Musser, J. et al. (2019). Environmental pH and peptide signaling control virulence of Streptococcus pyogenes via a quorum-sensing pathway. Nat Commun. 10, 2586.

Duranton, F., Cohen, G., De Smet, R., Rodriguez, M., Jankowski, J., Vanholder, R. et al. (2012). Normal and pathologic concentrations of uremic toxins. J. Am. Soc. Nephrol. 23, 1258–1270.

Eddy, S. (2001). Non-coding RNA genes and the modern RNA world.. Nat Rev Genet. 2, 919–929. doi: 10.1038/35103511.

Fricker, R., Brogli, R., Luidalepp, H., Wyss, L., Fasnacht, M., Joss, O. et al. (2019). A tRNA half modulates translation as stress response in Trypanosoma brucei. Nat Commun. 10, 118.

Fris, M. and Murphy, E. (2016). Riboregulators: Fine-Tuning Virulence in Shigella. Front Cell Infect Microbiol. 6, 2. doi: 10.3389/fcimb.2016.00002.

Furukawa, K., Ramesh, A., Zhou, Z., Weinberg, Z., Vallery, T., Winkler, W. et al. (2015). Bacterial riboswitches cooperatively bind Ni(2+) or Co(2+) ions and control expression of heavy metal transporters. Mol. Cell 57, 1088–1098.

García Véscovi, E., Soncini, F. and Groisman, E. (1996). Mg2+ as an extracellular signal: environmental regulation of Salmonella virulence. Cell 84, 165–174.

Gardner, P. P., Barquist, L., Bateman, A., Nawrocki, E. P., Weinberg, Z. (2011) RNIE: genome-wide prediction of bacterial intrinsic terminators. Nucleic Acids Res.39(14): 5845–5852.

Gollnick, P., Babitzke, P., Antson, A. and Yanofsky, C. (2005). Complexity in regulation of tryptophan biosynthesis in Bacillus subtilis. Annu. Rev. Genet. 39, 47–68.

Gripenland, J., Netterling, S., Loh, E., Tiensuu, T., Toledo-Arana, A. and Johansson, J. (2010). RNAs: regulators of bacterial virulence. Nat Rev Microbiol. 8, 857–866. doi: 10.1038/nrmicro2457.

Groisman, E., Cromie, M., Shi, Y. and Latifi, T. (2006). A Mg2+-responding RNA that controls the expression of a Mg2+ transporter. Cold Spring Harb Symp Quant Biol. 71, 251–258. doi: 10.1101/sqb.2006.71.005.

Groisman, E., Hollands, K., Kriner, M., Lee, E., Park, S. and Pontes, M. (2013). Bacterial Mg2+ homeostasis, transport, and virulence. Annu Rev Genet. 47, 625–646. doi: 10.1146/annurev-genet-051313-051025.

Grundy, F. and Henkin, T. (1993). tRNA as a positive regulator of transcription antitermination in B. subtilis. Cell 74, 475–482.

Guragain, M., King, M., Williamson, K., Perez-Osorio, A., Akiyama, T., Khanam, S. et al. (2016). The Pseudomonas aeruginosa PAO1 Two-Component Regulator CarSR Regulates Calcium Homeostasis and Calcium-Induced Virulence Factor Production through Its Regulatory Targets CarO and CarP. J. Bacteriol. 198, 951–963.

Guragain, M., Lenaburg, D., Moore, F., Reutlinger, I. and Patrauchan, M. (2013). Calcium homeostasis in Pseudomonas aeruginosa requires multiple transporters and modulates swarming motility. Cell Calcium 54, 350–361.

Ha, D. and O’Toole, G. (2015). c-di-GMP and its Effects on Biofilm Formation and Dispersion: a Pseudomonas Aeruginosa Review. Microbiol Spectr 3, 0003–2014.

Hacker, J. and Kaper, J. (2000). Pathogenicity islands and the evolution of microbes. Annu. Rev. Microbiol. 54, 641–679.

Hay, A., Yang, M., Xia, X., Liu, Z., Hammons, J., Fenical, W. et al. (2017). Calcium Enhances Bile Salt-Dependent Virulence Activation in Vibrio cholerae. Infect. Immun. 85,

Heroven, A., Nuss, A. and Dersch, P. (2017). RNA-based mechanisms of virulence control in Enterobacteriaceae. RNA Biol. 14, 471–487.

Hör, J., Gorski, S. and Vogel, J. (2018). Bacterial RNA Biology on a Genome Scale.. Mol Cell 70, 785–799. doi: 10.1016/j.molcel.2017.12.023.

Hughes, D. and Sperandio, V. (2008). Inter-kingdom signalling: communication between bacteria and their hosts.. Nat Rev Microbiol. 6, 111–120. doi: 10.1038/nrmicro1836.

Imazawa, R., Takahashi, Y., Aoki, W., Sano, M. and Ito, M. (2016). A novel type bacterial flagellar motor that can use divalent cations as a coupling ion.. Sci Rep. 6, 19773. doi: 10.1038/srep19773.

Johansson, J., Mandin, P., Renzoni, A., Chiaruttini, C., Springer, M. and Cossart, P. (2002). An RNA thermosensor controls expression of virulence genes in Listeria monocytogenes.. Cell 110, 551–561.

Juttukonda, L. and Skaar, E. (2015). Manganese homeostasis and utilization in pathogenic bacteria. Mol. Microbiol. 97, 216–228.

Kariisa, A., Weeks, K. and Tamayo, R. (2016). The RNA Domain Vc1 Regulates Downstream Gene Expression in Response to Cyclic Diguanylate in Vibrio cholerae. PLoS ONE 11, e0148478.

Kersey, C., Agyemang, P. and Dumenyo, C. (2012). CorA, the magnesium/nickel/cobalt transporter, affects virulence and extracellular enzyme production in the soft rot pathogen Pectobacterium carotovorum. Mol. Plant Pathol. 13, 58–71.

Kulkarni, P., Jia, T., Kuehne, S., Kerkering, T., Morris, E., Searle, M. et al. (2014). A sequence-based approach for prediction of CsrA/RsmA targets in bacteria with experimental validation in Pseudomonas aeruginosa. Nucleic Acids Res. 42, 6811–6825.

Lapouge, K., Schubert, M., Allain, F. and Haas, D. (2008). Gac/Rsm signal transduction pathway of gamma-proteobacteria: from RNA recognition to regulation of social behaviour. Mol. Microbiol. 67, 241–253.

Lebreton, A. and Cossart, P. (2017). RNA-and protein-mediated control of Listeria monocytogenes virulence gene expression. RNA Biol. 14, 460–470.

Leimeister-Wächterchter, M., Haffner, C., Domann, E., Goebel, W. and Chakraborty, T. (1990). Identification of a gene that positively regulates expression of listeriolysin, the major virulence factor of listeria monocytogenes.. Proc Natl Acad Sci U S A. 87, 8336–8340.

Leonard, S., Meyer, S., Lacour, S., Nasser, W., Hommais, F. and Reverchon, S. (2019). APERO: a genome-wide approach for identifying bacterial small RNAs from RNA-Seq data. Nucleic Acids Res.

Loh, E., Dussurget, O., Gripenland, J., Vaitkevicius, K., Tiensuu, T., Mandin, P. et al. (2009). A trans-acting riboswitch controls expression of the virulence regulator PrfA in Listeria monocytogenes. Cell 139, 770–779. doi: 10.1016/j.cell.2009.08.046.

Mao, C., Abraham, D., Wattam, A., Wilson, M., Shukla, M., Yoo, H. et al. (2015). Curation, integration and visualization of bacterial virulence factors in PATRIC. Bioinformatics 31, 252–258.

Massé E, Gottesman S. (2002) A small RNA regulates the expression of genes involved in iron metabolism in Escherichia coli. Proc Natl Acad Sci U S A. 99, 4620–4625.

Mastropasqua, M., Lamont, I., Martin, L., Reid, D., D’Orazio, M. and Battistoni, A. (2018). Efficient zinc uptake is critical for the ability of Pseudomonas aeruginosa to express virulence traits and colonize the human lung. J Trace Elem Med Biol. 48, 74–80.

Mengaud, J., Dramsi, S., Gouin, E., Vazquez-Boland, J., Milon, G. and Cossart, P. (1991). Pleiotropic control of Listeria monocytogenes virulence factors by a gene that is autoregulated.. Mol Microbiol. 5, 2273–2283.

Miyajima, A., Shibuya, M., Kuchino, Y. and Kaziro, Y. (1981). Transcription of the E. coli tufB gene: Cotranscription with four tRNA genes and inhibition by guanosine-5}-diphosphate-3}-diphosphate. Molecular and General Genetics MGG 183, 13–19. doi: 10.1007/BF00270131.

Morita, M., Tanaka, Y., Kodama, T., Kyogoku, Y., Yanagi, H. and Yura, T. (1999). Translational induction of heat shock transcription factor sigma32: evidence for a built-in RNA thermosensor.. Genes Dev 13, 655–665.

Mukai, T. (2021) Bioinformatic Prediction of an tRNASec Gene Nested inside an Elongation Factor SelB Gene in Alphaproteobacteria. Int. J. Mol. Sci. 22, 4605

Mulhbacher, J., Brouillette, E., Allard, M., Fortier, L., Malouin, F. and Lafontaine, D. (2010). Novel riboswitch ligand analogs as selective inhibitors of guanine-related metabolic pathways. PLoS Pathog. 6, e1000865.

Naghdi, M., Smail, K., Wang, J., Wade, F., Breaker, R. and Perreault, J. (2017). Search for 5’-leader regulatory RNA structures based on gene annotation aided by the RiboGap database. Methods 117, 3–13. doi: 10.1016/j.ymeth.2017.02.009.

Nahvi, A., Sudarsan, N., Ebert, M., Zou, X., Brown, K. and Breaker, R. (2002). Genetic control by a metabolite binding mRNA. Chem Biol 9, 1043.

Nelson, J., Sudarsan, N., Furukawa, K., Weinberg, Z., Wang, J. and Breaker, R. (2013). Riboswitches in eubacteria sense the second messenger c-di-AMP. Nat. Chem. Biol. 9, 834–839.

Nelson, J., Sudarsan, N., Phillips, G., Stav, S., Lunse, C., McCown, P. et al. (2015). Control of bacterial exoelectrogenesis by c-AMP-GMP. Proc. Natl. Acad. Sci. U.S.A. 112, 5389–5394.

Nies, D. (2019). The ancient alarmone ZTP and zinc homeostasis in Bacillus subtilis. Mol. Microbiol.

Nuss, A., Heroven, A. and Dersch, P. (2017). RNA Regulators: Formidable Modulators of Yersinia Virulence. Trends Microbiol. 25, 19–34.

Pacheco, A., Curtis, M., Ritchie, J., Munera, D., Waldor, M., Moreira, C. et al. (2012). Fucose sensing regulates bacterial intestinal colonization. Nature 492, 113–117. doi: 10.1038/nature11623.

Palmer, L. and Skaar, E. (2016). Transition Metals and Virulence in Bacteria. Annu Rev Genet. 50, 67–91. doi: 10.1146/annurev-genet-120215-035146.

Papp-Wallace, K. and Maguire, M. (2006). Manganese transport and the role of manganese in virulence. Annu. Rev. Microbiol. 60, 187–209.

Perreault, J., Weinberg, Z., Roth, A., Popescu, O., Chartrand, P., Ferbeyre, G. et al. (2011). Identification of hammerhead ribozymes in all domains of life reveals novel structural variations. PLoS Comput. Biol. 7, e1002031.

Price, I., Gaballa, A., Ding, F., Helmann, J. and Ke, A. (2015). Mn(2+)-sensing mechanisms of yybP-ykoY orphan riboswitches. Mol. Cell 57, 1110–1123.

Raina, M. and Ibba, M. (2014). tRNAs as regulators of biological processes. Front Genet. 5, 171.

Ramesh, A. and Winkler, W. (2010). Magnesium-sensing riboswitches in bacteria. RNA Biol. 7, 77–83.

Rasko, D., Moreira, C., Li, D., Reading, N., Ritchie, J., Waldor, M. et al. (2008). Targeting QseC signaling and virulence for antibiotic development. Science 321, 1078–1080. doi: 10.1126/science.1160354.

Remy, L., Carriere, M., Derre-Bobillot, A., Martini, C., Sanguinetti, M. and Borezee-Durant, E. (2013). The Staphylococcus aureus Opp1 ABC transporter imports nickel and cobalt in zinc-depleted conditions and contributes to virulence. Mol. Microbiol. 87, 730–743.

Ryckelynck, M., Giege, R. and Frugier, M. (2005). tRNAs and tRNA mimics as cornerstones of aminoacyl-tRNA synthetase regulations. Biochimie 87, 835–845.

Sarkisova, S., Lotlikar, S., Guragain, M., Kubat, R., Cloud, J., Franklin, M. et al. (2014). A Pseudomonas aeruginosa EF-hand protein, EfhP (PA4107), modulates stress responses and virulence at high calcium concentration. PLoS ONE 9, e98985.

Sarkisova, S., Patrauchan, M., Berglund, D., Nivens, D. and Franklin, M. (2005). Calcium-induced virulence factors associated with the extracellular matrix of mucoid Pseudomonas aeruginosa biofilms. J. Bacteriol. 187, 4327–4337.

Sayers, S., Li, L., Ong, E., Deng, S., Fu, G., Lin, Y. et al. (2019). Victors: a web-based knowledge base of virulence factors in human and animal pathogens. Nucleic Acids Res. 47, D693–D700.

Sharma, A., Dhasmana, N., Dubey, N., Kumar, N., Gangwal, A., Gupta, M. et al. (2017). Bacterial Virulence Factors: Secreted for Survival. Indian J. Microbiol. 57, 1–10.

Sherlock, M., Sudarsan, N. and Breaker, R. (2018). Riboswitches for the alarmone ppGpp expand the collection of RNA-based signaling systems. Proc. Natl. Acad. Sci. U.S.A. 115, 6052–6057.

Shi, Y., Zhao, G. and Kong, W. (2014). Genetic analysis of riboswitch-mediated transcriptional regulation responding to Mn2+ in Salmonella. J. Biol. Chem. 289, 11353–11366.

Skuzeski, J., Bozarth, C. and Dreher, T. (1996). The turnip yellow mosaic virus tRNA-like structure cannot be replaced by generic tRNA-like elements or by heterologous 3’ untranslated regions known to enhance mRNA expression and stability. J. Virol. 70, 2107–2115.

Stav, S., Atilho, R., Mirihana Arachchilage, G., Nguyen, G., Higgs, G. and Breaker, R. (2019). Genome-wide discovery of structured noncoding RNAs in bacteria. BMC Microbiol. 19, 66.

Sudarsan, N., Lee, E., Weinberg, Z., Moy, R., Kim, J., Link, K. et al. (2008). Riboswitches in eubacteria sense the second messenger cyclic di-GMP. Science 321, 411–413.

Tamayo, R. (2019). Cyclic diguanylate riboswitches control bacterial pathogenesis mechanisms. PLoS Pathog. 15, e1007529.

Tang, Q., Yin, K., Qian, H., Zhao, Y., Wang, W., Chou, S. et al. (2016). Cyclic di-GMP contributes to adaption and virulence of Bacillus thuringiensis through a riboswitch-regulated collagen adhesion protein. Sci Rep. 6, 28807.

Vakulskas, C., Potts, A., Babitzke, P., Ahmer, B. and Romeo, T. (2015). Regulation of bacterial virulence by Csr (Rsm) systems. Microbiol. Mol. Biol. Rev. 79, 193–224.

Valentini, M. and Filloux, A. (2016). Biofilms and Cyclic di-GMP (c-di-GMP) Signaling: Lessons from Pseudomonas aeruginosa and Other Bacteria. J. Biol. Chem. 291, 12547–12555.

Valverde, C., Lindell, M., Wagner, E. and Haas, D. (2004). A repeated GGA motif is critical for the activity and stability of the riboregulator RsmY of Pseudomonas fluorescens. J. Biol. Chem. 279, 25066–25074.

Velasco, E., Wang, S., Sanet, M., Fernandez-Vazquez, J., Jove, D., Glaria, E. et al. (2018). A new role for Zinc limitation in bacterial pathogenicity: modulation of Î±-hemolysin from uropathogenic Escherichia coli. Sci Rep. 8, 6535.

Véscovi, E., Ayala, Y., Di Cera, E. and Groisman, E. (1997). Characterization of the bacterial sensor protein PhoQ. Evidence for distinct binding sites for Mg2+ and Ca2+. J Biol Chem. 272, 1440–1443.

Wattam, A., Abraham, D., Dalay, O., Disz, T., Driscoll, T., Gabbard, J. et al. (2014). PATRIC, the bacterial bioinformatics database and analysis resource. Nucleic Acids Res. 42, D581–D591. doi: 10.1093/nar/gkt1099.

Wedekind, J., Dutta, D., Belashov, I. and Jenkins, J. (2017). Metalloriboswitches: RNA-based inorganic ion sensors that regulate genes. J. Biol. Chem. 292, 9441–9450.

Weinberg, Z., Lunse, C., Corbino, K., Ames, T., Nelson, J., Roth, A. et al. (2017). Detection of 224 candidate structured RNAs by comparative analysis of specific subsets of intergenic regions. Nucleic Acids Res. 45, 10811–10823.

Weinberg, Z., Wang, J., Bogue, J., Yang, J., Corbino, K., Moy, R. et al. (2010). Comparative genomics reveals 104 candidate structured RNAs from bacteria, archaea, and their metagenomes. Genome Biol. 11, R31.

Wishart, D., Feunang, Y., Marcu, A., Guo, A., Liang, K., Vazquez-Fresno, R. et al. (2018). HMDB 4.0: the human metabolome database for 2018. Nucleic Acids Res. 46, D608–D617.

Wishart, D., Jewison, T., Guo, A., Wilson, M., Knox, C., Liu, Y. et al. (2013). HMDB 3.0--The Human Metabolome Database in 2013. Nucleic Acids Res. 41, D801–807.

Wishart, D., Knox, C., Guo, A., Eisner, R., Young, N., Gautam, B. et al. (2009). HMDB: a knowledgebase for the human metabolome. Nucleic Acids Res. 37, D603–610.

Wishart, D., Tzur, D., Knox, C., Eisner, R., Guo, A., Young, N. et al. (2007). HMDB: the Human Metabolome Database. Nucleic Acids Res. 35, D521–526.

Zeenko, V., Ryabova, L., Spirin, A., Rothnie, H., Hess, D., Browning, K. et al. (2002). Eukaryotic elongation factor 1A interacts with the upstream pseudoknot domain in the 3’ untranslated region of tobacco mosaic virus RNA. J. Virol. 76, 5678–5691.

Zhang, X., Liu, Z., Yi, J., Tang, H., Xing, J., Yu, M. et al. (2012). The tRNA methyltransferase NSun2 stabilizes p16INK4 mRNA by methylating the 3’-untranslated region of p16. Nat Commun. 3, 712.

